# Pancreatic cancer risk predicted from disease trajectories using deep learning

**DOI:** 10.1101/2021.06.27.449937

**Authors:** Davide Placido, Bo Yuan, Jessica X. Hjaltelin, Chunlei Zheng, Amalie D. Haue, Piotr J Chmura, Chen Yuan, Jihye Kim, Renato Umeton, Gregory Antell, Alexander Chowdhury, Alexandra Franz, Lauren Brais, Elizabeth Andrews, Debora S. Marks, Aviv Regev, Siamack Ayandeh, Mary Brophy, Nhan Do, Peter Kraft, Brian M. Wolpin, Nathanael Fillmore, Michael Rosenthal, Søren Brunak, Chris Sander

**Author notes:** joint first authors (Davide Placido, Bo Yuan, Jessica X. Hjaltelin, Chunlei Zheng). joint senior authors (Søren Brunak, Chris Sander).

## Abstract

Pancreatic cancer is an aggressive disease that typically presents late with poor patient outcomes. There is a pronounced medical need for early detection of pancreatic cancer, which can be addressed by identifying high-risk populations. Here we apply artificial intelligence (AI) methods to a dataset of 6 million patient records with 24,000 pancreatic cancer cases in the Danish National Patient Registry (DNPR) and, for comparison, a dataset of three million records with 3,900 pancreatic cancer cases in the United States Department of Veterans Affairs (US-VA) healthcare system. In contrast to existing methods that do not use temporal information, we explicitly train machine learning models on the time sequence of diseases in patient clinical histories and test the ability to predict cancer occurrence in time intervals of 3 to 60 months after risk assessment.

For cancer occurrence within 36 months, the performance of the best model (AUROC=0.88, DNPR), trained and tested on disease trajectories, exceeds that of a model without longitudinal information (AUROC=0.85, DNPR). Performance decreases when disease events within a 3 month window before cancer diagnosis are excluded from training (AUROC[3m]=0.83). Independent training and testing on the US-VA dataset reaches comparable performance (AUROC=0.78, AUROC[3m]=0.76). These results raise the state-of-the-art level of performance of cancer risk prediction on real-world data sets and provide support for the design of prediction-surveillance programs based on risk assessment in a large population followed by affordable surveillance of a relatively small number of patients at highest risk. Use of AI on real-world clinical records has the potential to shift focus from treatment of late-stage to early-stage cancer, benefiting patients by improving lifespan and quality of life.

## Introduction

### Clinical need for early detection

Pancreatic cancer is a leading cause of cancer-related deaths worldwide with increasing incidence (Rahib et al. 2014). Early diagnosis of pancreatic cancer is a key challenge, as the disease is typically detected at a late stage. Approximately 80% of pancreatic cancer patients are diagnosed with locally advanced or distant metastatic disease, when long-term survival is extremely uncommon (2-9% of patients at 5-years) (McGuigan et al. 2018). However, patients who present with early-stage disease can be cured by a combination of surgery, chemotherapy and radiotherapy. Indeed, more than 80% of patients with stage IA pancreatic ductal adenocarcinoma (PDAC) achieve 5-year overall survival [National Cancer Institute, USA, (Blackford et al. 2020)]. Thus, a better understanding of the risk factors for pancreatic cancer and detection at early stages has great potential to improve patient survival and reduce overall mortality.

### Known risk factors of limited use

The incidence rate of pancreatic cancer is substantially lower compared with other high mortality cancers, such as lung, breast and colorectal cancer. Thus, age-based population screening is difficult due to poor positive predictive values for potential screening tests and large numbers of futile evaluations for patients with false-positive results. Moreover, few high-penetrance risk factors are known for pancreatic cancer impeding early diagnosis of this disease. Risk of pancreatic cancer has been assessed for many years based on family history, behavioral and clinical risk factors and, more recently, circulating biomarkers and genetic predisposition (Amundadottir et al. 2009; Petersen et al. 2010; D. Li et al. 2012; Wolpin et al. 2014; Klein et al. 2018; Kim et al. 2020). Currently, some patients with high risk due to family history or rare inherited pathogenic variants or cystic lesions of the pancreas undergo serial pancreas-directed imaging to detect early pancreatic cancers, but these patients account for less than 20% of those who develop pancreatic cancer.

To address the challenge of early detection of pancreatic cancer in the general population (Pereira et al. 2020; Singhi et al. 2019), we aim to predict the risk of pancreatic cancer from real-world longitudinal clinical records and identify high-risk patients, which will facilitate the design of surveillance programs for early detection. Real-world data are usually noisy and the predictive power can be affected by various confounding factors, therefore, the development of realistic risk prediction methods requires a choice of appropriate machine learning methods, in particular deep learning techniques that work on large and **noisy** sequential datasets (Dietterich 2002; LeCun, Bengio, and Hinton 2015).

### Earlier clinical ML work

We build on earlier work in the field of risk assessment based on clinical data and disease trajectories using machine learning technology (Nielsen et al. 2019; Thorsen-Meyer et al. 2020). AI methods have been successfully applied to a number of clinical decision support problems (Shickel et al. 2018), such as choosing optimal time intervals for actions in intensive care units (Hyland et al. 2020), assessing cancer risk from images (Esteva et al. 2017; Yala et al. 2019; Yamada et al. 2019), predicting the risk of potentially acute disease progression, such as in kidney injury (Tomašev et al. 2019) and the likelihood of a next diagnosis based on past EHR sequences, in analogy to natural language processing (Y. Li et al. 2020; Thorsen-Meyer et al. 2022).

### Earlier ML work on PDAC risk

For risk assessment of pancreatic cancer, recently machine learning predictive models using patient records have been built using health interview survey data (Muhammad et al. 2019), general practitioners’ health records controlled against patients with other cancer types (Malhotra et al. 2021), real-world hospital system data (Appelbaum, Cambronero, et al. 2021; X. Li et al. 2020), and from an electronic health record (EHR) database provided by TriNetX, LLC. (Chen et al. 2021; Appelbaum, Berg, et al. 2021). While demonstrating the information value of health records for cancer risk, these previous studies used only the occurrence of disease codes, not the time sequence of disease states in a patient trajectory - in analogy to the ‘bag-of-words’ models in natural language processing that ignore the actual sequence of words. Previous studies had used the Danish health registries to generate population-wide disease trajectories, but in a non-predictive manner (Hu et al. 2019; Jensen et al. 2014).

### Advance here - better data and better ML

Here we exploit the power of advanced machine learning technology by focusing on the time sequence of clinical events and by predicting the risk of cancer occurrence in specific time intervals after assessment. This investigation was first carried out using the Danish National Patient Registry (DNPR), which contains data from hospital admissions including out-patient visits from 1977 to 2018 (41 years), including diagnosis codes, for a total of 8.6 million patients. About 40,000 of these had a diagnosis of pancreatic cancer (Schmidt et al. 2015; Siggaard et al. 2020). To maximize predictive information extraction from these records we tested a range of machine learning methods. These methods range from regression methods and machine learning without time dependence to time series methods such as Gated Recurrent Units (GRU) and Transformer, adapting AI methods that have been very successful in natural language processing and analysis of other time series data (Cho et al. 2014; Tealab 2018; Vaswani et al. 2017).

### Advance - prediction time intervals

The likely action resulting from a personalized positive prediction of cancer risk ideally should take into account the probability of the disease occurring within a shorter or longer time frame. For this reason, we designed the prediction method to predict not only whether cancer is likely to occur, but also to provide risk assessment in incremental time intervals following the assessment, where time of assessment is defined as the day on which the risk prediction is performed based on the history of clinical records of the particular patient. We also analyzed which diagnoses in a patient’s history of diagnosis codes are most informative of cancer risk - not as isolated factors but in the context of the person’s complete history of disease codes. Finally, we propose a practical scenario for surveillance programs, taking into consideration the availability of real-world data, the accuracy of prediction on such data, the scope of a surveillance program, the likely cost and success rate of surveillance methods and the overall potential benefit of early treatment (Supplement, **Result S1**).

## Results

### Datasets

#### Dataset of disease trajectories: Denmark-DNPR

We used disease trajectory data from the Danish National Patient Registry (DNPR). Demographic information was obtained by linkage to the Central Person Registry, which is possible via the personal identification number introduced in 1968 and uniquely identifies any Danish citizen over the lifespan (Schmidt, Pedersen, and Sørensen 2014). DNPR covers approximately 8.6 million patients with 229 million hospital diagnoses, with on average 26.7 diagnosis codes per patient. For training we used trajectories of ICD diagnostic codes with explicit time stamps for each hospital contact comprising diagnoses down to the three-character category in the ICD hierarchy. The accuracy of ICD codes for pancreatic cancer, which were used to label cases in training, was estimated to be above 88% (Methods). We used data from January 1977 to April 2018 and filtered out patients with discontinuous or very short trajectories (<5 events in total), ending up with 6.2 million patients (**Figure S1A**). The case cohort includes 23,985 pancreatic cancer (PC) cases with cancer diagnosed at a median age of 70 years (49.5% man, 50.5 female) (**Figure 2**, **Table S1**).

**Figure 1.**
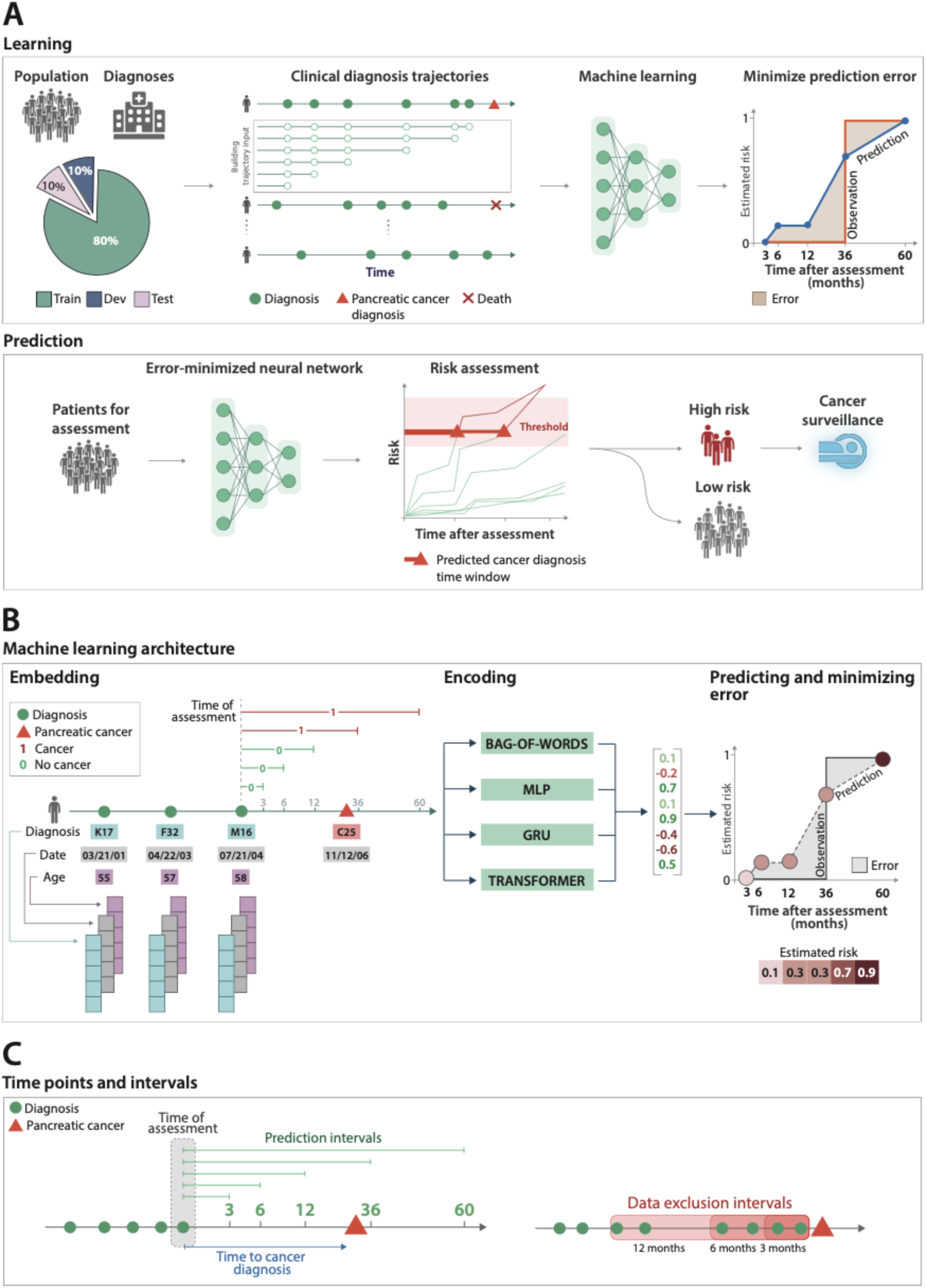
Training and prediction of pancreatic cancer risk from disease trajectories. (**A**) Learning: The general machine learning workflow starts with partitioning the data into training set (Train), development set (Dev) and test set (Test). The trajectories for training input are generated by sampling continuous subsequences of diagnoses for each patient’s diagnosis history, each starting with the first record but with different end points. The training and development sets are used for training machine learning models to fit a risk score function (prediction) to a step function (observation) that represents the occurrence of a pancreatic cancer diagnosis, by minimizing the prediction error over all instances. Prediction: A model’s ability to accurately predict is evaluated using the withheld ‘test’ set. The prediction model, depending on the prediction threshold selected from among possible operational points, discriminates between patients at higher and lower risk of pancreatic cancer. The risk model can guide the development of surveillance initiatives. (**B**) The model trained with real-world clinical data has three steps: embedding, encoding and prediction. The embedding machine transforms categorical disease codes and time stamps of these disease codes into a lower-dimensional real-number continuous space. The encoding machine extracts information from a disease history and summarizes each sequence in a characteristic fingerprint in the latent space (vertical vector). The prediction machine then uses the fingerprint to generate predictions for cancer occurrence within different time intervals after the time of assessment (3, 6, 12, 36, 60 months). The model parameters are trained by minimizing the difference between the predicted and the observed cancer occurrence. (**C**) Terminology for time points and intervals. The end point of a disease trajectory is the time of assessment. From the time of assessment, cancer risk is assessed within 3, 6, 12, 36 and 60 months. To test the influence of close-to-cancer diagnosis codes on the prediction of cancer occurrence, exclusion intervals are used to remove diagnoses in the last 3, 6 and 12 months before cancer diagnosis.

**Figure 2.**
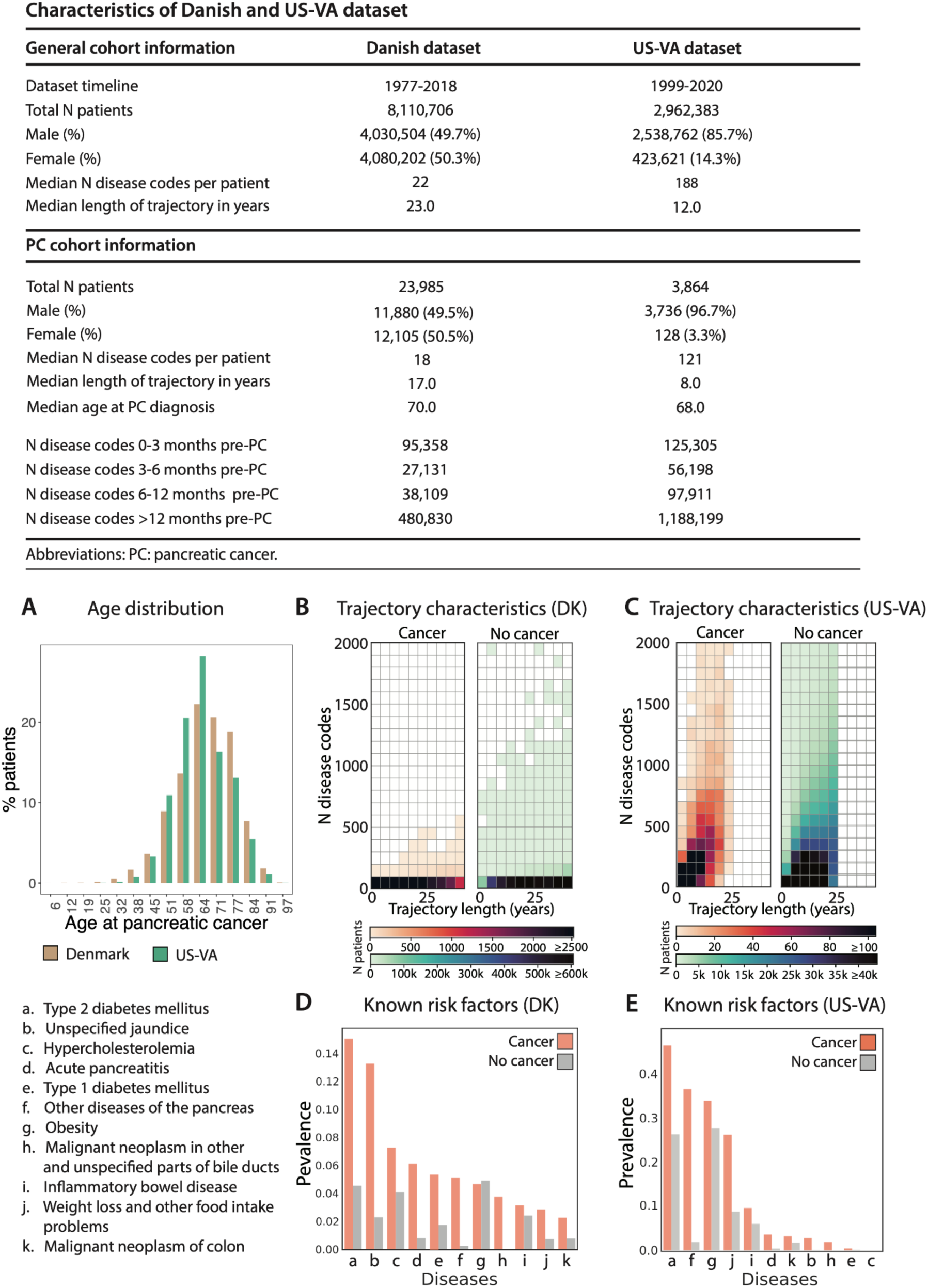
Characteristics of the Danish (DK) and US-Veterans Administration (VA) patient registries. (**A**) Age distributions for age at pancreatic cancer diagnosis in the two cohorts. (**B,C**) The DK dataset (DNPR) has a longer median length of disease trajectories, but lower median number of disease codes per patient compared to the VA dataset, so the machine learning process, independently in each dataset, has to cope with very different distributions of disease trajectories in terms of length of trajectories and density of the number of disease codes. Color level indicates the number of patients in a given bin. (**D,E**) Background check on the distribution disease code in the clinical records: prevalence of known risk factors in cancer vs. non-cancer patients in the DK (**D**) and VA (**E**) datasets, counting whether a disease code occurred at least once in a patient’s history previous to their pancreatic cancer code (cancer) or two years previous to the end of data (no cancer).

#### Dataset of disease trajectories: US-VA

For validation in another health care system, we used clinical records from the United States Department of Veterans Affairs (US-VA) hospital system via the VA Corporate Data Warehouse, which integrates both electronic health record and cancer registry data from VA facilities nationwide. As in the Danish dataset, we also used explicit longitudinal records from US-VA, i.e., trajectories of ICD codes with explicit time stamps.

Cases of pancreatic cancer were ascertained using VA cancer registry data. The accuracy of the US-VA cancer registry data for pancreatic cancer, which was used to label cases in training, was estimated to be 95-96% (Methods). We used data from 1999 to 2020 and filtered out patients with less than five recorded diagnosis codes (**Figure S1b, Figure 2**). The selected dataset has 3.0 million patients with 3,864 pancreatic cancer patients (**Figure S1b**). On average, the health records in the US- VA dataset have shorter (median 12 years) but substantially denser disease history (median 188 records per patient). For US-VA, the median age of pancreatic cancer diagnosis is 68.0 years (96.7% male, 3.3% female). These numerical differences likely reflect the differences in population (entire population in Denmark vs military veterans in US-VA), in the health care systems in the two datasets, such as referral, billing and documentation practices.

### Model architecture

#### Network architecture/layers

The machine learning model for predicting cancer risk from disease trajectories consists of: (1) **input:** data for each event in a trajectory (diagnosis code and time stamps), (2) **embedding:** the event features onto real number vectors, (3) **encoding:** the trajectories in a lower-dimensional latent space, and (4) **predicting:** time-dependent cancer risk. (1) **Input**: In order to best exploit the longitudinality of the health/disease records and provide an opportunity to discover early indicators of cancer risk, all contiguous subsequences of diagnoses from a patient’s history were sampled, starting with the earliest record and ending at a series of time points, with increasing gaps between the end of the trajectory and cancer occurrence for positive cases (Methods). The partial trajectories provide information in support of prediction for different time spans between risk assessment and cancer occurrence, rather than just binary prediction that cancer will occur at any time after assessment. (2) **Embedding**: Each item in a disease trajectory is an event denoted with a level-3 ICD code. To extract informative features from such high-dimensional input, the machine learning process is designed to embed the categorical input vectors into a continuous, lower-dimensional space. Temporal information, i.e. diagnosis dates and age at each diagnosis are also embedded (Methods). The mapping of the input to the embedding layer is trained together with other parts of the model. (3) **Encoding**: The longitudinal nature of the disease trajectories allows us to construct time-sequence models using sequential neural networks, such as gated recurrent units (GRU) models (Cho et al. 2014). We also used the Transformer model (Vaswani et al. 2017) which uses an attention mechanism to capture complex sequential interdependencies. For comparison, we also tested a bag-of-words (i.e., bag-of-disease-codes) approach that ignores the time and order of disease events by pooling the elements of the event vectors. (4) **Predicting:** The encoded latent space vector is then used as input to a multilayer fully-connected network to estimate the risk of cancer within distinct prediction intervals ending a few months or several years after the end of a trajectory (the time of risk assessment).

#### Prediction of occurrence within a time interval and data exclusion

For each of the disease trajectories ending at time *t_a_*, a 5-dimensional risk score is calculated, where each dimension represents the cumulative risk of cancer occurrence within a particular prediction window after *t_a_*, e.g., ending at *t_a_* + 12 months (Lin et al. 2008; Yala et al. 2021). The risk score is constrained to monotonically increase with time as the risk of cancer occurrence naturally increases over time, for a given disease trajectory. If and when the risk score exceeds a prediction threshold, cancer diagnosis is predicted to have occurred (**Figure 1**). In this way, the model uses a time sequence of disease codes for one person as input and separately predicts a cancer diagnosis to occur within 3, 6, 12, 36, or 60 months after the time *t_a_* of risk assessment. In addition, to test the influence of close-to-cancer diagnosis codes on the prediction of cancer occurrence, exclusion intervals are used to remove diagnoses in the last 3, 6 and 12 months before cancer diagnosis during training.

#### Scanning hyperparameters for each model type

To comprehensively test the performance of different types of machine learning models, we first conducted an extensive search over hyperparameters and selected the best set of hyperparameters for each model, and then selected the best model type. The model types included transformer, GRU, a multilayer perceptron (MLP) and bag-of-words. Each model was tested on specific hyperparameter configurations (**Table S2**). To avoid overfitting and to test generalizability of model predictions, we partitioned patient records randomly into 80%/10%/10% training/development/test sets. We conducted training only on the training set and used the development set to examine the performance for different hyperparameter settings, which guides model selection. Subsequently, the performance of the selected models was evaluated on the fully withheld test set and reported as an estimate of performance in prospective patients from the health system used to train and test the models.

### Evaluation of model performance

#### Picking a best model - DK

We evaluated the different models trained in the Danish DNPR using the AUROC and relative risk (RR) curves (**Figure 3A-H**). All performance metrics are calculated on the basis of applying each trained risk assessment model to the test set. The test set is strictly withheld during training and hyperparameter search. In the final performance evaluation of different types of machine learning models on the test sets, the models which explicitly use and encode the time sequence of disease codes, i.e., GRU and Transformer, ranked highest by AUROC (**Figure 3A; Table S3**). For the prediction of cancer incidence within 3 years of the assessment date (the date of risk prediction), the Transformer model had the best performance (AUROC=0.879 [0.877-0.880]), followed by GRU (AUROC=0.852 [0.850-0.854]). In order to gain a better intuition regarding the impact of applying a model in a real case scenario, we also report the relative risk score (RR) of cancer patients in the high-risk group as predicted by the machine learning model.

**Figure 3.**
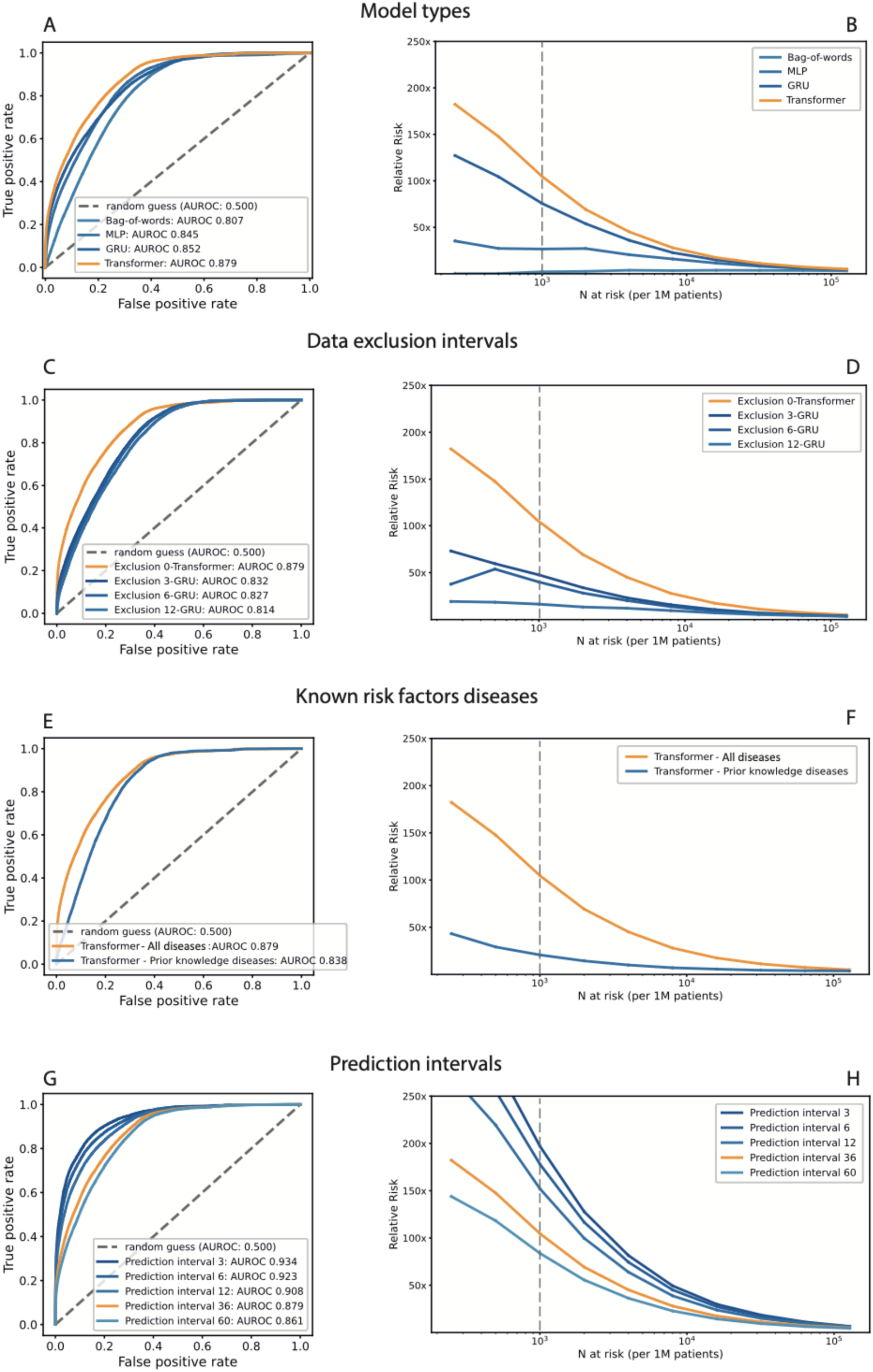
Performance of the machine learning (ML) on clinical record trajectories in predicting pancreatic cancer occurrence, in the DK data set. For each model and prediction evaluation, performance is better for larger AUROC (**A,C,E,G**) and for higher relative risk for the N highest-risk patients (cumulative; **B,D,F,H**). **(A,B)** Choice of algorithm: the transformer algorithm is best with AUROC=0.879 (no data exclusion, 36 months prediction interval). **(C,D)** Choice of input data: Prediction performance declines with exclusion, in training, of k=3,6,12 months of data between the end of a disease trajectory and cancer occurrence (best model for each exclusion interval, for 36 month prediction interval). **(E,F)** Choice of input data: Prediction is better for all 2000 ICD level-3 disease codes used throughout in training (Methods) compared to only the subset of 23 known risk factors; using transformer, all data, for 36 month prediction interval. **(G,H)** Choice of prediction task: Prediction of cancer is more difficult for larger prediction intervals, the time interval within which cancer is predicted to occur after assessment (transformer model, all data). We evaluate prediction performance for the 36-month prediction interval (orange) in the other panels, as this is of practical interest. **(B,D,F,H)** Prediction performance at a **particular operational point**, e.g., N=1000 highest-risk patients out of 1 million, the relative risk (RR) is 104.7 for the 36 month prediction interval using all data and 47.6 with 3 month data exclusion.

The relative risk score is defined at a given operational decision point (Methods). This assesses by what factor a prediction method does better than a random model (**Figure 3B**). The relative risk (RR) for the 36 months prediction interval is 104.7 for the Transformer model (with time sequences), at an operational point defined by the N=1000 highest risk patients out of 1 million patients (0.1% at highest risk; notation: N1000).

### 3A Comparison with previous models

Earlier work also developed machine learning methods on real-world clinical records to predict pancreatic cancer risk (Appelbaum, Cambronero, et al. 2021; Appelbaum, Berg, et al. 2021; Chen et al. 2021; X. Li et al. 2020). These previous studies had encouraging results, but neither used the time sequence of disease histories to extract time-sequential longitudinal features. For comparison we implemented analogous approaches, a bag-of-words model and an MLP model. We evaluated the non-time-sequential models on the DNPR dataset, and the performance for predicting cancer occurrence within 36 months was AUROC=0.807 (0.805-0.809) for the bag-of-words model and 0.845 (0.843-0.847) for the MLP model (**Figure 3A**). The RR is also much lower (2.1 and 26.6, respectively), compared to that of time-sequential models (e.g., 104.7 for Transformer).

### 3C Performance with data exclusion

Disease codes within a very short time before diagnosis of pancreatic cancer are most probably directly indicative such that even without any machine learning, well-trained clinicians would include pancreatic cancer in their differential diagnosis. Disease codes just prior to pancreatic cancer occurrence may also encompass it (e.g., neoplasm of the digestive tract) and thus directly reflect the label one wants to infer. To avoid undue influence of these disease codes in training, we therefore separately trained the models excluding input diseases diagnoses from the last 3/6/12 months prior to the diagnosis of pancreatic cancer (‘data exclusion’). As expected, when training with data exclusion, the performance decreased from AUROC=0.879 to AUROC=0.843/0.829/0.827 for 3/6/12 months data for the best model - all for prediction of cancer occurrence within 36 months (DNPR dataset, **Figure 3C**).

### 3E Performance for training on known risk factors rather than a full ICD set

One can train on a smaller or larger set of disease codes that occur in patient disease trajectories. E.g., one can use prior knowledge and limit the input for training to known risk factors, i.e., diseases that have been reported to be indicative of the likely occurrence of pancreatic cancer (Yuan et al. 2020; Klein 2021). We find that prediction performance with the subset of ICD codes for 23 known risk factors reduces prediction accuracy of the transformer model to AUROC=0.838 compared to 0.879 for all diagnosis codes, and therefore we used the latter throughout the rest of the work (**Figure 3E,F, Table S4**).

### 3G Prediction for time intervals

It is of particular clinical interest to consider the risk of cancer over different time intervals. The machine learning models in this work are designed to report risk scores for pancreatic cancer occurrence within 3, 6, 12, 36 and 60 months of the date of risk assessment. As expected, it is more challenging to predict cancer occurrence within longer rather than shorter time intervals, as the longer time intervals allow for larger time gaps between the end of the disease trajectory (time of assessment) and the time of cancer diagnosis. Indeed, prediction performance for the transformer model decreases from an AUROC of 0.908 (0.906-0.911) for cancer occurrence within 12 months to an AUROC of 0.879 (0.877-0.880) for occurrence within 3 years (without data exclusion) (**Figure 3G,H**).

#### Information contribution as a function of time gap between of assessment and cancer occurrence

The exclusion of trajectories ending very close to pancreatic cancer removes the influence of disease codes that represent late symptoms of pancreatic cancer or are otherwise easily attributable to pancreatic cancer. However, data exclusion of such late events alone does not quantify the influence of longer term risk factors on prediction. We therefore computed the recall rate of prediction as a function of the time-to-cancer, defined as the time between the end of disease trajectory and the occurrence of cancer (**Figure 5A, C**). As expected, recall levels decrease with longer time-to-cancer, from 8% for cancer occurring about 1 year after assessment to a recall of 4% for cancer occurring about 3 years after assessment (DNPR, **Figure 5A**). This suggests that the model not only learns from symptoms very close to pancreatic cancer but also from longer disease history, albeit at lower accuracy.

**Figure 4.**
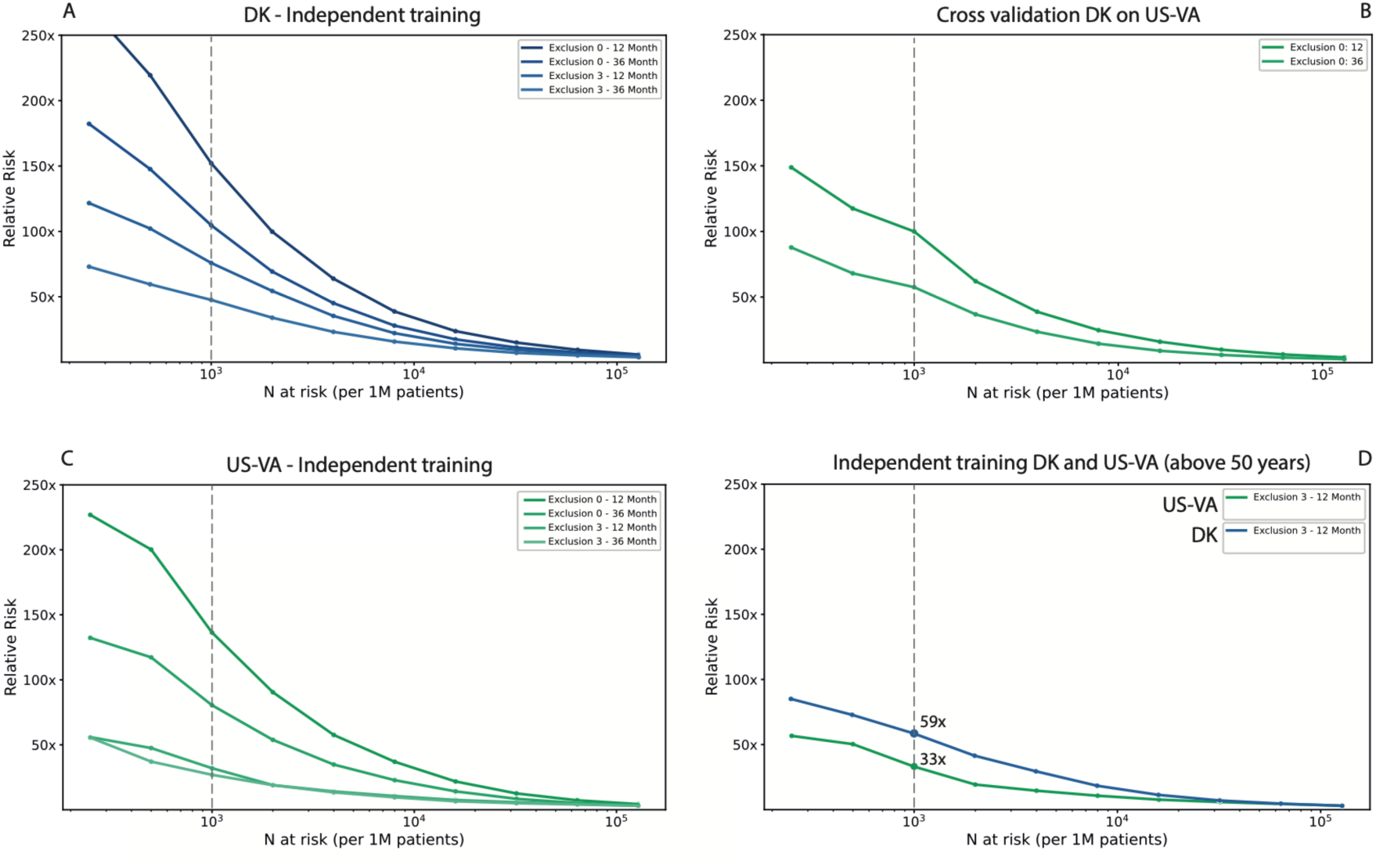
Estimated performance of a surveillance program for high risk patients in different health systems and with different operational choices. Estimated relative risk for the top N high-risk patients is based on computing the accuracy of prediction on the withheld test set **(A,C,D)** or external dataset **(B)**. In designing surveillance programs, one has a choice between models trained on all data (excl=0) versus models trained excluding data from the last three months before cancer occurrence (excl=3) and between prediction for cancer within 12 months of prediction or within 36 months, with estimates of prospective performance based on independent training on data from the DK **(A)** Performance for the US-VA **(C)** health systems. **(B)** Estimated performance is somewhat lower for cross-application for a model trained on DK data applied to VA patient data, with an AUROC of 0.710 (0.708-0.712), illustrating the challenge of deriving globally valid prediction tools without independent localized or system-specific training. **(D)** A proposed practical choice for a surveillance program with good estimated accuracy of prediction, in either system, would entail application of independently trained models with 3-month data exclusion for a prediction interval of 12 months for patients above age 50.

**Figure 5.**
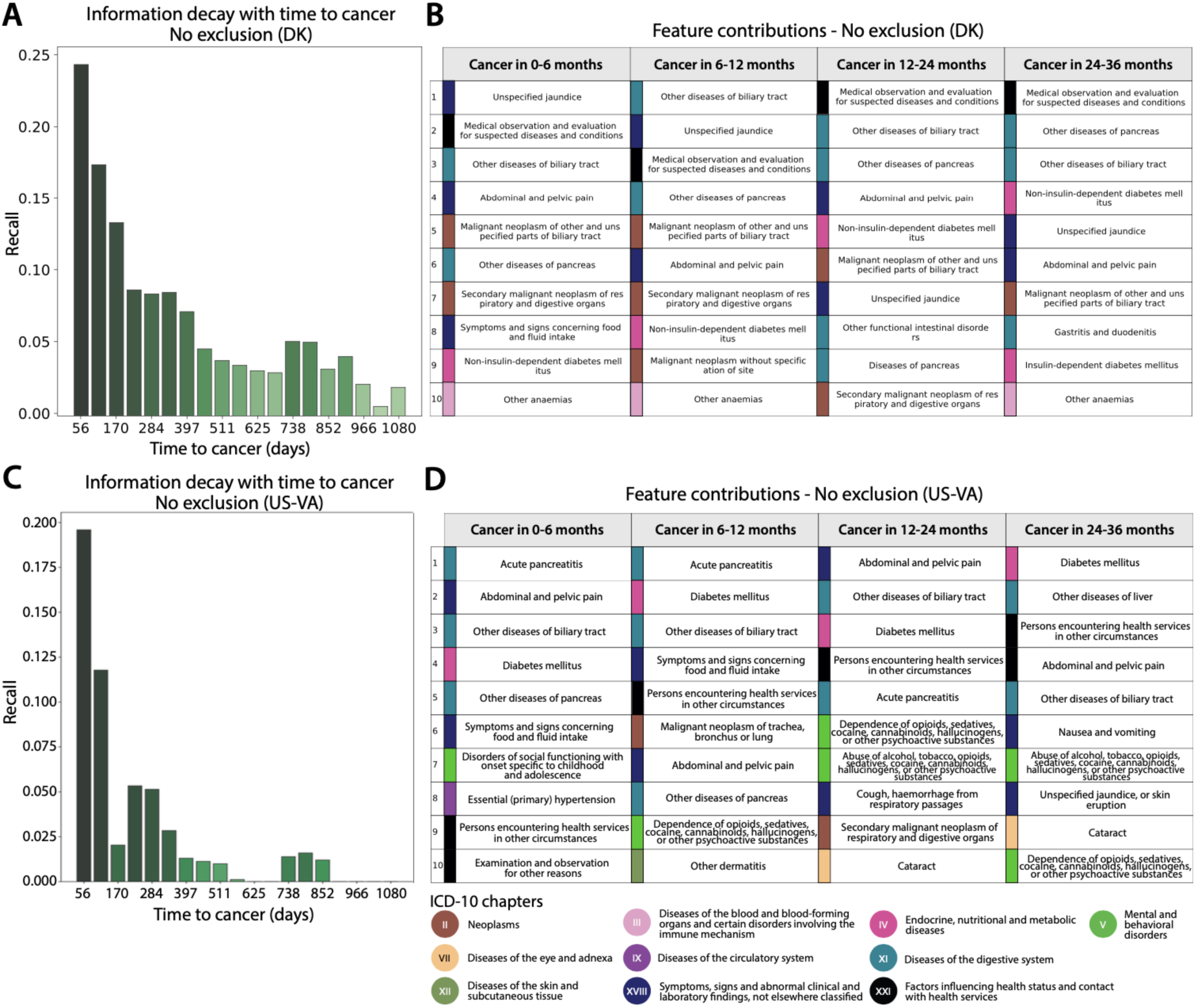
Predictive capacity and feature contributions of disease trajectories. (**A,C**) Distribution of recall (sensitivity) values at the F1 operational point (Methods) as a function of time-to-cancer (time between the end of a disease trajectory and cancer diagnosis). The recall values decrease with time-to-cancer. (**A**) Danish system, for models trained on all data (no exclusion). The contribution of age for DNPR is in Figure S3. (**C**) US-VA system, for models trained on all data. (**B,D**) Top 10 features that contribute to the cancer prediction in time-to-cancer intervals of 0-6, 6-12, 12-24 and 24-36 months for the **(B)** Danish and **(D)** US-VA systems. The features are sorted by the contribution score (**Suppl. Tables S5**). We used an integrated gradient (IG) method to calculate the contribution score for each input feature for each trajectory, then summed over all trajectories with cancer diagnosis within the indicated time interval.

### Generalization for different health care systems

#### 4B Performance by cross-application of a model to different datasets

In order to assess the predictive performance of the model in other health care systems, we applied the best machine learning model trained on the Danish dataset to disease trajectories of patients in the US-VA dataset, without any adaptation except for mapping the ICD codes from one system to the other. Prediction performance for cancer occurrence within 36 months after assessment declined from an AUROC of 0.879 (0.877-0.880) and RR=104.6, for a Denmark-trained transformer model applied to Danish DNPR patient data (test set), to an AUROC of 0.710 (0.708-0.712) and RR=57.4, for the same model applied to US-VA patient data (**Figure 4B**). The most striking difference in input data between the two systems is the shorter and more dense disease history in the US-VA trajectories compared to the Danish ones (**Figure 2B-C**).

#### 4C Performance by independent training on a different dataset

Motivated by the decrease in performance when testing the Denmark-derived model on the US-VA dataset, we trained and evaluated the model on the US-VA dataset from scratch. For the independently trained model, the performance is clearly higher than in cross-application, with a test-set AUROC of 0.775 (0.772-0.778) and RR=80.4 at the N1000 operational point (0.1% at highest risk) for cancer occurring within 36 months (**Figure 4C**). The difference in performance of the independently trained models in the two health care systems may be in part due to differences in medical and reporting practices or in demographics, including different age and sex distributions.

#### 4D Proposal for a realistic surveillance program

A realistic surveillance program has to use an operational point (decision threshold) taken into consideration cost and benefit in real-world clinical practice. While we have no empirical data on cost and benefit in this study, we can evaluate predictive performance, which is a key factor in assessing benefit. An example (**Figure 4D**), for a potentially realistic choice of model and decision threshold, is a surveillance program at an operational point of N=1000 (0.1%) highest risk patients out of 1 million patients, i.e., at a risk fraction of 0.1% of all patients assessed. At this point, the best independently trained models trained with 3 months of data exclusion evaluated for the 12 month prediction interval for patients above age 50 obtain a relative risk of RR=58.7 for DNPR (GRU model) and of RR=33.1 for US-VA (Transformer model). Corresponding models using all the data (no exclusion) have higher values of RR. Considerations for the design of a surveillance program are further discussed below.

### Predictive Features

#### Interpretation: contributing factors from gradient method

Although the principal criterion for the potential impact of prediction-surveillance programs is robust predictive performance, it is of interest to interpret the features of any predictive method: which diagnoses are most informative of cancer risk? Computational methods can infer the contribution of a particular input variable to the prediction by a machine learning engine, e.g., the integrated gradients (IG) algorithm (Sundararajan, Taly, and Yan 2017) (**Figure 5C-D; Table S5**). The IG contribution was calculated separately for different times between assessment and cancer diagnosis, in particular at 0-6 months, 6-12 months, 12-24 months and 24-36 months after assessment, for all patients who developed cancer. The aim was to explore how diagnosis codes contribute differently to the risk of pancreatic cancer, depending on how close to pancreatic cancer they were administered. As expected, there is a tendency for codes, which in normal clinical practice are known to indicate the potential presence of pancreatic cancer, to have a higher contribution to prediction for trajectories that end closer to cancer diagnosis. On the other hand, putative early risk factors have a higher IG score for trajectories that end many months before cancer diagnosis. With the proviso that the IG scores separately evaluate individual features that actually act collectively, these results provide some level of confidence in the machine learning prediction method. The interpretation of individual risk factors from the machine learning feature list as causative may be subject to misinterpretation as their contribution here is only evaluated in the context of complete disease histories. However, our main goal in this report is to achieve robust predictive power from disease trajectories, rather than mechanistic interpretations.

#### Interpretation: contributing early factors

The top contributing features extracted from the trajectories with time to cancer diagnosis in 0-6 months - such as unspecified jaundice, diseases of biliary tract, abdominal-pelvic pain, weight loss and neoplasms of digestive organs *-* may be symptoms of or otherwise closely related to pancreatic cancer (**Table S5A**). It is also of interest to identify early risk factors for pancreatic cancer. For trajectories with longer time between assessment and cancer diagnosis, other disease codes - such as type 2 diabetes and insulin-independent diabetes - make an increasingly large contribution, consistent with epidemiological studies (Yuan et al. 2020; Klein et al. 2013; Kim et al. 2020) and the observed disease distribution in the DNPR and US-VA datasets (**Figure 4, S3**). Other factors, such as cholelithiasis (gallstones) and reflux disease, are perhaps of interest in terms of potential mechanistic hypotheses, such as inflammation of the pancreas prior to cancer as a result of cholelithiasis or a hypothetical link between medication by proton pump inhibitors such as omeprazole in reflux disease and the effect of increased levels of gastrin on the state of the pancreas (Alkhushaym et al. 2020).

#### Interpretation: extent of similarity of IG features in the two systems

We have additionally performed the disease contribution analysis on the US-VA dataset (**Table S5B**). The disease contribution analysis showed commonly identified disease codes between the two systems such as unspecified jaundice, other disorders of pancreas, other diseases of biliary tract, diabetes mellitus, abdominal and pelvic pain, other diseases of liver, symptoms and signs concerning food and fluid intake. Generally, in addition to disease codes that are rather specific for pancreatic cancer, others are unspecific, indicating the importance of including many variables with limited individual prognostic value to obtain good prognostication.

## Discussion

### Advances in this work

We present a new framework for applying deep learning to longitudinal datasets of disease trajectories to predict the risk of a low-incidence but very aggressive cancer. This study was designed to make explicit use of the time sequence of disease events; and, to assess the ability to predict cancer risk for increasing intervals between the end of the disease trajectory and cancer occurrence. Earlier work has demonstrated the potential of applying AI methods to assess pancreatic cancer risk, but did not exploit the information in the temporal sequence of diseases (Appelbaum, Cambronero, et al. 2021; Chen et al. 2021). Our results indicate that using the time ordering in disease histories as input to the learning engine, rather than just disease occurrence at any time, significantly improves the predictive power of AI methods in anticipating pancreatic cancer occurrence.

### Performance in a different healthcare systems

A single, globally robust model that predicts cancer risk for patients in different countries and different healthcare systems remains elusive. Cross-application of the Danish model to the US-VA database had lower performance in spite of common use of ICD disease codes **(Figure 4)**. Reasons for this lack of transferability plausibly are differences in clinical practice, such as frequency of reporting disease codes in clinical records, typical thresholds for seeking medical attention, potential influence of billing constraints and billing optimization, as well as entry to and departure from the US-VA health system - all of these in contrast to the more uniform and comprehensive national nature of the Danish disease registries. However, the AI methods used are sufficiently robust to achieve a higher level of performance in the US-VA system via independent training, as opposed to cross-application of an externally trained prediction tool. We conclude that when there are significant differences in healthcare systems, independent model training in different geographical locations is required to achieve locally optimal model performance. However, if independent training is not feasible, e.g. in smaller health care systems, then cross-application may be of value, depending on the actual outcome of cross-performance tests, which can be performed on smaller datasets than those required for training.

### Application in clinical practice - 3 steps

Successful implementation of early diagnosis and treatment of pancreatic cancer in clinical practice will likely require three essential steps: (i) identification of high-risk patients, (ii) detection of early cancer or precancerous states by detailed surveillance of high-risk patients, and (iii) effective treatment after early detection (Singhi et al. 2019; Kenner et al. 2021). The overall impact in clinical practice depends on the success rates in each of these stages. This work only addresses the first stage. With a reasonably accurate method for predicting cancer risk one can direct appropriate high-risk patients into surveillance programs. A sufficiently enriched pool of high-risk patients would make detailed screening tests affordable, as such tests are likely to be prohibitively expensive at a population level, and enhance the positive predictive value of such tests.

### Design of surveillance programs

Based on the prediction accuracy reported here, one can design a potentially realistic prediction-surveillance process, in which software is applied to health records of, e.g., 1 million patients, followed by identification of those at highest risk and recruitment into a surveillance program with detailed screening tests for, e.g., 1000 highest-risk patients (0.1% of all assessed). Implementation requires choosing an operational point along the curve of relative risk against the fraction of patients at high risk (**Figures 3,4**) (or an equivalent point on a precision-recall curve) with an achievable high positive predictive value so as to reduce false positives in the high-risk group and therefore minimize unnecessary effort, potential harm from unnecessary invasive procedures and potential anxiety.

One proposal for the design of a surveillance program is to focus on patients from age 50 (**Figure 4D**). (1) One mode of operation is to supplement existing surveillance practice in a given hospital system, in which patients with, e.g., pancreatitis, abdominal pain and/or unspecified jaundice are routinely under surveillance by clinicians. To supplement these, using a model *trained with 3 months data exclusion* and applied to patients of age 50 or older, our results indicate that out of 1000 patients at highest risk (0.1% of 1 million) 70 patients would be future pancreatic cancer patients (PPV 0.07, 12 months prediction interval, GRU, DNPR), assuming that these typically are not already under surveillance. (2) Another mode of operation is to consider the risk prediction for any patient, including those that are already under surveillance as part of current clinical practice. In this mode one identifies patients at highest risk using a model *trained on all data* (no data exclusion), for which we computed a PPV of 0.32 (12 months prediction interval, Transformer, DNPR, age 50 or older), corresponding to 320 future pancreatic cancer patients out of 1000 at highest risk, (in a population of 1 million assessed by the prediction method). In addition to patients already under surveillance this would plausibly add a non-negligible number of additional candidates for surveillance.

Interestingly, the relative risk as reported here is comparable to that reported for the - surveillance of patients with genetic risk factors such as mutations in the BRCA2, ATM, PALB2 and other genes or those with a high value of a polygenic risk score (Klein 2021; Galeotti et al. 2021). While those with family history of pancreatic cancer are reported to have a 9x higher risk of pancreatic cancer, heritability is estimated to explain only 4-5% of all pancreatic cancers. Here, complementary to established surveillance programs for patients with elevated risk based on germline variants or family history, the focus is on the application to the entire patient population, for which genetic or detailed family history data is available only in a relatively small number of patients. The real-world AI approach casts a much wider net and is a practical approach in that running the prediction program routinely on, say, 1 million patients - is relatively inexpensive. In addition to prediction performance, cost of screening is then a leading factor in determining the fraction of high-risk patients nominated for a screening program in any particular health care system.

### Estimated performance of surveillance programs

In either scenario we estimate the pool of patients already under surveillance plus additional surveillance candidates would result in approximately 320 (out of 1000) patients (0.1% of 1 million), who actually will develop pancreatic cancer. Whether or not cancer is actually detected early in these patients of course depends on the clinical tests performed. Our study cannot provide estimates on the positive early detection rate of such a surveillance program, only on the number of patients estimated to eventually develop pancreatic cancer. Detailed deliberations with clinicians will be required to design the details of such a program, e.g., to choose the frequency of follow-up visits and tests to be performed, optimizing success rates of tests, minimizing cost and minimizing potential harm from invasive procedures. For cost considerations alone, we provide a hypothetical tradeoff algorithm (**Results S1**). A surveillance program will also have to be supplemented with counseling, in analogy to genetic counseling, to provide patients with an informed choice and to minimize potential anxiety. The numerical estimates of performance of the risk assessment tool presented here are simply one of several considerations needed for the design of a surveillance program.

In our view, the performance reported here, in particular in the Danish system, is an advance relative to earlier methods and is approaching a level in which the design of a surveillance program is a practical proposition. Even a relatively moderate level of additional early detection may be considered of value, provided that cost and implementation issues can be successfully addressed in a real-world implementation.

### Challenges for future improvements

We expect further increases in prediction accuracy with the real-world availability of data beyond disease codes, such as medication, laboratory values, observations in clinical notes, abdominal imaging (computerized tomography, magnetic resonance imaging), treatment records from general practitioners (Malhotra et al. 2021; Lemanska et al. 2022) as well as germline genetic profiles. To achieve a globally useful set of prediction rules, access to large data sets of disease histories aggregated nationally or internationally will be extremely valuable, with careful assessment of the accuracy of clinical records. An ideal scenario for a multi-institutional collaboration would be to employ federated learning across a number of different healthcare systems (Konečný et al. 2016). Federated learning obviates the need for sharing primary data and only requires permission to run logically identical computer codes at each location and then share and aggregate results. The challenges to achieve federated learning are, however, not only technical but also social and organizational, especially in a competitive health care landscape.

### Impact on patients

The particular advantage of a real-world high-risk prediction-surveillance process is that computational screening of a large population in the first step is inexpensive while the costly second step of sophisticated clinical screening and intervention programs is limited to a much smaller number of patients, those at highest risk. Prediction performance at the level shown here may be sufficient for an initial design of real world clinical prediction-surveillance programs and future improvements are likely. AI on real-world clinical records has the potential to produce a scalable workflow for early detection of cancer in the community, to shift focus from treatment of late- to early-stage cancer, improve the quality of life of patients, and increase the benefit/cost ratio of cancer care.

## Methods

### Processing of disease code trajectory datasets

#### The population-level Danish DNPR dataset

The first part of the project was conducted using a dataset of disease histories from the Danish National Patient Registry (DNPR), covering all 229 million hospital diagnoses of 8.6 million patients between 1977-2018. This includes inpatient contacts since 1977 and outpatient and emergency department contacts since 1995, but not data from the general practitioners’ records (Schmidt et al. 2015). DNPR access was approved by the Danish Health Data Authority (FSEID-00003092 and FSEID-00004491.) Each entry of the database includes data on the start and end date of an admission or visit, as well as diagnosis codes. The diagnoses are coded according to the International Classification of Diseases (ICD-8 until 1994 and ICD-10 since then). The accuracy of cancer diagnosis disease codes, as examined by the Institute of Clinical Medicine, Aarhus University Hospital, has been reported to be 98% (accuracy for 19 Charlson conditions in 950 reviewed records) (Thygesen et al. 2011). For cancer diagnoses specifically, the reference evaluation was based on detailed comparisons between randomly sampled discharges from five different hospitals and review of a total of 950 samples (Schmidt et al. 2015). We used both the ICD-8 code 157 and ICD-10 code C25, *malignant neoplasm of pancreas*, to define pancreatic cancer (PC) cases.

#### Use of ICD codes

The ICD classification system has a hierarchical structure, from the most general level, e.g., *C: Neoplasms*, to the most specific four-character subcategories e.g. *C25.1: Malignant neoplasm of body of pancreas*. The Danish version of the ICD-10 is more detailed than the international ICD-10 but less detailed than the clinical modification of the ICD-10 (ICD-10-CM). In this study, we used the three-character category ICD codes (n=2,997) in constructing the predictive models and defined “pancreatic cancer (PC) patients” as patients with at least one code under *C25: Malignant neoplasm of pancreas*. For the diagnosis codes in the DNPR, we removed disease codes labeled as ‘temporary’ or ‘referral’ (8.3% removed, **Figure S1A**), as these can be misinterpreted when mixed with the main diagnoses and are not valuable for the purposes of this study.

#### Filtering the Danish dataset

Danish citizens have since 1968 been assigned a unique lifetime Central Person Registration (CPR) Number, which is useful for linking to person-specific demographic data. Using these we retrieved patient status as to whether patients are active or inactive in the CPR system as well as information related to residence status. We applied a demographic continuity filter. For example, we excluded from consideration residents of Greenland, patients who lack a stable place of residence in Denmark, as these would potentially have discontinuous disease trajectories. By observation time we mean active use of the healthcare system.

#### Subset of Danish dataset used for training

The Danish DNPR dataset comprised a total of 8,110,706 patients, of which 23,601 had the ICD-10 pancreatic cancer code *C25* and 14,720 had the ICD-8 pancreatic cancer code *157*. We used both ICD-10 and ICD-8 independently, without semantic mapping, while retaining the pancreatic cancer occurrence label, assuming that machine learning is able to combine information from both. Subsequently, we removed patients that have too few diagnoses (<5 events). The number of positive patients used for training after applying the length filter are 23,985 (82% ICD-10 and 18% ICD-8). Coincidentally, this resulted in a more strict filtering for ICD-8 events which were used only in 1977-1994. The final dataset was then randomly split into training (80%), development (10%) and test (10%) data, with the condition that all trajectories from a patient were only included in one split group (train/dev/test), to avoid any information leakage between training and development/test datasets.

#### Quality control using the Danish Cancer Registry dataset

To perform a quality check of the pancreatic cancer cases in DNPR, we compared pancreatic cancer cases from both DNPR with the pancreatic cancer cases in the Danish Cancer Registry (DCR). DCR is a population-wide cancer cohort and one of the largest and most comprehensive in the world, which has recorded nation-wide cancer incidences from 1943 (Gjerstorff 2011; Sundhedsstyrelsen 2009). The DCR contains information on diagnosis, cancer type, tumor topography, TNM staging classification and morphology. The purpose of DCR is to keep track of cancer incidences and mortality in Denmark and to study causes, courses and statistics for treatment improvements in the Danish health system. The cancer registry administration was moved from The Danish Cancer Society to the National Board of Health in 1997.

We compared the overlap between the pancreatic cancer cases in DNPR versus the cases in DCR to estimate the quality of our case population used for training. 88% of the pancreatic cancer cases used for training overlapped with the cancer registry. Due to the differing nature of the DCR and DNPR, for which the latter is a more administrative registry with multiple coding purposes such as hospital reimbursement, monitoring hospital services and patient trajectories, quality assurance etc., we did not expect these registries to overlap 100%. As there may be correctly labeled cases in DNPR that did not get captured in DCR, the 88% concordance is a lower bound on accuracy of the pancreatic cancer ICD-10 codes in DNPR, which we used as case labels in training.

#### The military veterans US-VA dataset

Analysis in the US-VA system was conducted with approval from the VA Boston Healthcare System Institutional Review Board. For the US-VA system, we used data from the VA Corporate Data Warehouse (CDW), which collates electronic health record and cancer registry information on veterans treated at VA facilities nationwide (Price, Shea, and Gephart 2015), using methods similar to prior work (Elbers et al. 2020; Chang et al. 2022)(Wu et al. 2022). The CDW includes electronic health record data from 1999 to the present originating from the comprehensive range of primary care and specialized services the VA provides at its inpatient and outpatient facilities, as well as claims data for care received at outside facilities and paid for by the VA. Each outpatient visit in the database is associated with the date it occurred, and each inpatient visit is associated with an admission and discharge date. Both inpatient and outpatient visits are also associated with International Classification of Diseases codes (ICD-9 before October 1, 2015 and ICD-10 on and after that date) pertaining to the visit.

#### The US-VA cancer registry and data quality

In addition to electronic health record data, the CDW also includes cancer registry data. VA cancer registry activities were initiated pursuant to a national directive in 1998, with incident cases annotated retrospectively from 1995 and prospectively until the present (Zullig et al. 2012). Cases are abstracted manually by trained cancer registrars in accordance with standards of the North American Association of Central Cancer Registries (M. L. Thornton 2022). Potential cases are flagged for review by custom software (OncoTraX) which identifies potential cases automatically based on occurrence of structured data such as ICD codes in the electronic health record. This approach, where semi-automated screening is followed by manual review by trained cancer registrars, results in highly accurate case ascertainment (Zullig et al. 2012, 2019). For example, one study found that VA cancer registry data in the CDW had near perfect accuracy in colorectal cancer case ascertainment by all evaluated measures (positive predictive value, negative predictive value, sensitivity, and specificity) as compared to de novo manual review of 200 potential cases (Earles et al. 2018). In contrast, case ascertainment using ICD codes from the electronic health record had only 58% positive predictive value. Regarding pancreatic cancer case ascertainment specifically, we compared pancreatic cancer cases in the VA cancer registry to cases identified based on ICD codes in the VA electronic health record. We found that 94.9% of pancreatic cases in the VA cancer registry between 1999 and 2020 had at least one ICD-10 code for pancreatic cancer in the electronic health record, while only 32.4% of patients with a pancreatic cancer ICD code in the electronic health record were found in the cancer registry. Spurious ICD codes for pancreatic cancer can occur in the EHR due to multiple reasons, including outright error, use of the cancer code for a screening or evaluation visit, use of the cancer code for medical history, and use of the cancer code in dual-use scenarios where cancer care is provided outside the VA.

#### Filtering the US-VA dataset and subset used for training

VA patients were included in the study as described in the flow chart (**Figure S1B**). Out of all 15,933,326 patients with ≥1 ICD code in US-VA Corporate Data Warehouse between 1999 and 2020 and available in our research study database, we randomly sampled a subset of approximately 3 million patients (2,975,110) due to limitations of the computational resources available. Based on the considerations above, we identified patients with pancreatic cancer as the overlap of those with a diagnosis of pancreatic cancer in the VA cancer registry and those with an ICD code in the VA electronic health record system. We excluded patients with an ICD code for pancreatic cancer in the VA electronic health record data who did not have an entry for pancreatic cancer in the VA cancer registry or vice versa, since the status of these patients was unclear. We further excluded patients with short trajectories (<5 visits), as in the Danish dataset. This resulted in a final VA dataset of 1,948,209 patients total, including 3,418 patients with pancreatic cancer. We randomly allocated patients in the final VA dataset into training (80%), development (10%), and test (10%) sets, with the condition that all trajectories from a patient were only included in one split group (train/dev/test), to avoid any information leakage between training and development/test datasets, as in the Danish dataset.

#### Similarity of survival curves in the two health care systems

As a check on the quality of pancreatic cancer case ascertainment in the Danish and US-VA datasets, we plotted overall survival in each dataset, stratified by cancer stage (**Figure S1C)**. Cancer stage was obtained from the respective dataset’s cancer registry. Stage was only available on a subset of patients. The overall similarity of the survival curves in the two datasets adds confidence to the quality of the data, as selected for the comparative study.

### Training

The following processing steps were carried out identically for DNPR and US-VA datasets. For each patient, whether or not they ever had pancreatic cancer, the data was augmented by using all continuous **partial trajectories** of (minimal length >=5 diagnoses) from the beginning of their disease history and ending at different time points, which we call the time of assessment. For cancer patients, we used only trajectories that end before cancer diagnoses, i.e. t_a_<t_cancer_<t_death_. We used a **step function annotation** indicating cancer occurrence at different time points (3, 6, 12, 36, 60 months) after the end of each partial trajectory. For the positive (‘PC’) cases this provides the opportunity to learn from disease histories with a significant time gap between the time of assessment and the time of cancer occurrence. For example, for a patient, who had pancreatitis a month or two just before the cancer diagnosis, it is of interest to learn which earlier disease codes might have been predictive of cancer occurrence going back at least several months or perhaps years. The latter is also explored by separately re-training of the machine learning model excluding data from the last three or six months before cancer diagnosis.

For patients without a pancreatic cancer diagnosis we only include trajectories that end earlier than 2 years before the end of their disease records (due to death or the freeze date of the DNPR data used here). This avoids the uncertainty of cases in which undiagnosed cancer might have existed before the end of the records. The datasets were sampled in small batches for efficient computation, as is customary in machine learning. Due to the small number of cases of pancreatic cancer compared to controls, we used balanced sampling from the trajectories of the patients in the training set such that each batch has an approximately equal number of positive (cancer) and negative (non-cancer) trajectories.

### Model development

A desired model for such diagnosis trajectories consists of three parts: embedding of the categorical disease features, encoding time sequence information, and assessing the risk of cancer. We embed the discrete and high-dimensional disease vectors in a continuous and low-dimensional latent space (Mikolov et al. 2013; Gehring et al. 2017). Such embedding is data-driven and trained together with other parts of the model. For machine learning models not using embedding, each categorical disease was represented in numeric form as a one-hot encoded vector. The longitudinal records of diagnoses allowed us to construct time-sequence models with sequential neural networks. After embedding, each sequence of diagnoses, was encoded into a feature vector using different types of sequential layers (recurrent neural network, RNN, and gated recurrent units, GRU), attention layers (transformer), or simple pooling layers (bag-of-words model and multilayer perceptron model [MLP]). The encoding layer also included age inputs, which has been demonstrated to have a strong association with pancreatic cancer incidence (Klein 2021). Finally, the embedding and encoding layers were connected to a fully-connected feedforward network (FF) to make predictions of future cancer occurrence following a given disease history (the bag-of-words model only uses a single linear layer).

The model output consists of a risk score that monotonically increases for each time interval in the follow-up period after risk assessment. As cancer by definition occurs before cancer diagnosis, the risk score at a time point *t* is interpreted as quantifying the risk of cancer occurrence between *t_a_*, the end of the disease trajectory (the time of assessment), and time *t = t_a_* + 3, 6, 12, 36, 60 months. For a given prediction threshold, scores that exceed such threshold at time *t* are considered to indicate cancer occurrence prior to *t*. We currently do not distinguish between different stages of cancer, neither in training from cancer diagnoses nor in the prediction of cancer occurrence.

The model parameters were trained by minimizing the prediction error quantified as the difference between the observed cancer diagnosis in the form of a step function (0 before the occurrence of cancer, 1 from the time of cancer diagnosis) and the predicted risk score in terms of a positive function that monotonically increases from 0, using a cross-entropy loss function, with the sum over the five time points, and L2 regularization on the parameters (**Figure 1A**).

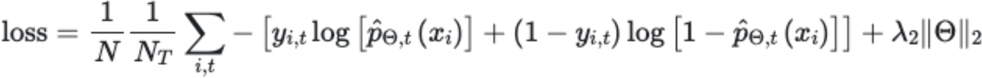

where *t* ∈ {3,6,12,36,60} months; *N_T_* = 5 for non-cancer patient and *N_T_* ≤ 5 for cancer patients where we only use the time points before the cancer diagnosis; *i* = 1,2,3, …, *N* labels samples; *Θ* is the set of model parameters; *λ*_2_ is the regularization strength; *p̂* is the estimated risk output by the model; *x_i_* are the input disease trajectories, *y_i,t_* = 1 for cancer occurrence and *y_i,t_* = 0 for no cancer within *t*-month time window.

The transformer model, unlike the recurrent models, does not process the input as a sequence of time steps but rather uses an attention mechanism to enhance the embedding vectors correlated with the outcome. In order to enable the transformer to digest temporal information such as the order of the exact dates of the diseases inside the sequence, we used positional embedding to encode the temporal information into vectors which were then used as weights for each disease token. Here we adapted the positional embedding from (Vaswani et al. 2017) using the values taken by cosine waveforms at 128 frequencies observed at different times. The times used to extract the wave values were the age at which each diagnosis was administered and the time difference between each diagnosis. In this way the model is enabled to distinguish between the same disease assigned at different times as well as two different disease diagnoses far and close in time. The parameters in the embedding machine, which addresses the issue of data representation suitable for input into a deep learning network, were trained together with the encoding and prediction parts of the model with back propagation (**Figure 2**).

To comprehensively test different types of neural networks and the corresponding hyperparameters, we conducted a large parameter search for each of the network types (**Table S2**). The different types of models include simple feed-forward models (LR, MLP) and more complex models that can take the sequential information of disease ordering into consideration (GRU and Transformer). See supplementary table with comparison metrics across different models (**Table S3**). In order to estimate the uncertainty of the performances, the 95% confidence interval was constructed using 200 resamples of bootstrapping with replacement.

### Evaluation

The evaluation was carried out separately for each prediction interval of 0-3, 0-6, 0-12, 0-36, and 0-60 months. For example, consider the prediction score for a particular trajectory at the end of the 3 year prediction interval (**Figure1C**). If the score is above the threshold, one has a correct positive prediction, if cancer has occurred at any time within 3 years; and a false positive prediction, if cancer has not occurred within 3 years. If the score is below the threshold, one has a false negative prediction if cancer has occurred at any time within 3 years; and a true negative prediction, if cancer has not occurred within 3 years. As both training and evaluation make use of multiple trajectories per patient, with different end-of-trajectory points, the performance numbers are computed over trajectories.

The relative risk ratio (RR) is calculated as the odds of getting pancreatic cancer when classified at high risk compared to a random method that just uses the disease incidence in the population. RR is defined as

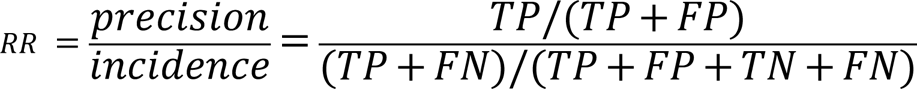

where TP = True Positives, FP = False Positives, FN = False Negatives, TN = True negatives. The relative risk score is defined at a given operational decision point along the RR curve as a function of the number of patients predicted to be at high risk **(Figure 3 right; Figure 4)**. The relative risk ratio assesses by what factor a prediction method does better than a random pick based just on the population disease incidence.

For the purpose of comparing different algorithms, input modes and DK-DNPR vs. US-VA, we report exhaustive performance tables (**Supplement Tables S3A-I**) with precision and recall at the F1 operational point, which maximizes the harmonic mean of recall and precision (Sasaki 2007). However, for consideration of clinical implementation, which requires severe limits on the number of patients that can be advanced to a surveillance program, we use an operational point for the top 1000 high risk patients out of 1 million patients (0.1% of 1 million, see relative risk plots **Figures 3&4**).

### Cross-application to the US-VA dataset

In order to assess the predictive performance of the model in other health care systems, we applied the best machine learning model trained on the Danish DNPR to disease trajectories of patients in a US Veterans Administration dataset. For the US-VA dataset, we directly applied models without any adaptation except for mapping the ICD code from one system to the other. In brief, ICD-9 codes in the US-VA dataset were first mapped to ICD-10 codes, followed by adding a prefix “D” to obtain Danmark compatible ICD-10 codes.

### Interpreting clinically relevant features

In order to find the features that are strongly associated with pancreatic cancer, we have used an attribution method for neural networks called integrated gradients (Sundararajan, Taly, and Yan 2017). This method calculates the contribution of input features, called attribution, cumulating the gradients calculated along all the points in the path from the input to the baseline. We chose the output of interest to be the 36-month prediction. Positive and negative attribution scores (contribution to prediction) indicate positive correlation with pancreatic cancer patients and non-pancreatic-cancer patients, respectively. Since the gradient cannot be calculated with respect to the indices used as input of the embedding layer, the input used for the attribution was the output of the embedding layer. Then, the attribution at the token level was obtained summing up over each embedding dimension and summing across all the patient trajectories. Similarly, for each trajectory, we calculated the age contribution as the sum attribution of the integrated gradients of the input at the age embedding layer.

### Software

The software will be made freely available in source code on a github repository.

## Acknowledgements

We thank Adam Yala for expert advice and contributions to methods and software, Regina Barzilay for discussions and guidance, Barbara Kenner for advice, and Julia Sidenius and Inna Chen for advice on clinical matters and known risk factor diseases. DP, JXH, ADH and SB acknowledge support from the Novo Nordisk Foundation (grants NNF17OC0027594 and NNF14CC0001). BMW acknowledges support from the Hale Family Center for Pancreatic Cancer Research, Lustgarten Foundation Dedicated Laboratory program, Stand Up to Cancer, NIH grant U01 CA210171, NIH grant P50 CA127003, Pancreatic Cancer Action Network, Noble Effort Fund, Wexler Family Fund, Promises for Purple, and Bob Parsons Fund. MHR acknowledges support from the Hale Family Center for Pancreatic Cancer Research, Lustgarten Foundation Dedicated Laboratory program, and NIH grant U01 CA210171. We thank Stand Up to Cancer (grant SU2C#6180), the Lustgarten Foundation and their donors for financial and community support. The authors thank Shawn Murphy and Henry Chueh and the Mass General Brigham Health Care Research Patient Data Registry group for facilitating use of their database.

## Supplementary Materials

**Figure S1.** Preprocessing and filtering of the DK and US-VA disease trajectory datasets. Filtering of the Danish (DK) and US-VA patient registries prior to training. In the Danish dataset, patient status codes were used to remove discontinuous disease histories such as patients living in Greenland, patients with alterations in their patient ID or patients who lack a stable residence in Denmark. We also removed referral and temporary diagnosis codes which are not the final diagnosis codes and can be misleading to use for training. Patients with short trajectories (<5 diagnosis codes) were removed. The final set of patients were split into Training (80 %), Validation (10%) and Testing set (10%). For the US-VA dataset, around 3 million of patients were randomly sampled due to computational limitations and patients with ICD-9/10 for pancreatic cancer, but without entries in US-VA cancer registry were excluded. Similar to the Danish dataset filtering, short trajectories (<5 diagnosis codes) were removed and patients were split into Training (80 %), Validation (10%) and Test set (10%).

**Figure S1A.**
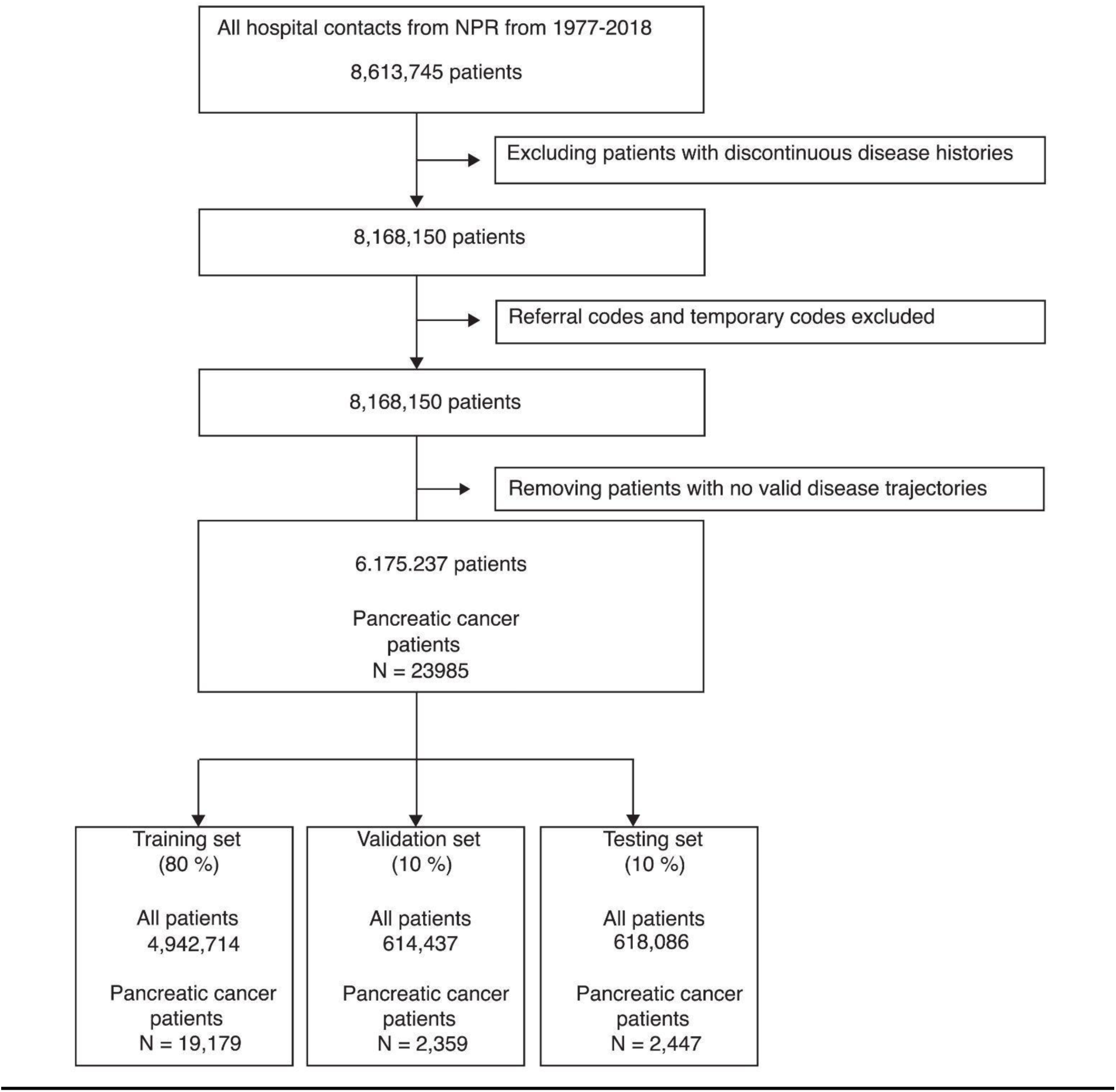
Denmark (DK) DNPR.

**Figure S1B.**
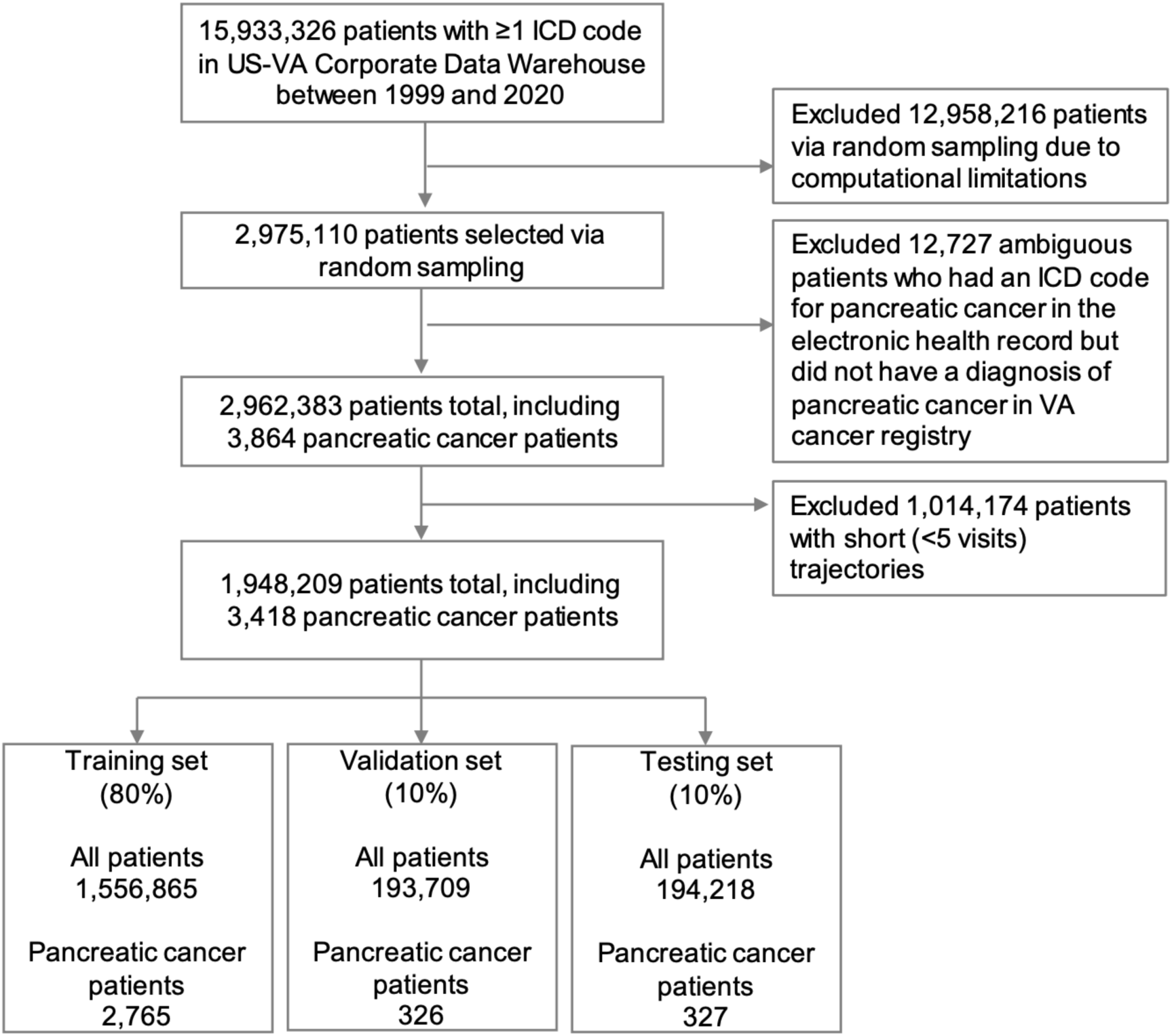
US-VA.

**Figure S1C.** Survival curves.

Overall survival in each dataset, stratified by cancer stage. Cases were ascertained using the methods described in Methods, and cancer stage was obtained from the respective dataset’s cancer registry. Stage was only available on a subset of patients.

For the Danish dataset, five-year survival was 23% for Stage I, 8.3% for Stage II, 2.3% for Stage III, and 0.8 for Stage IV. Median survival was 645 days for Stage I, 483 days for Stage II, 262 days for Stage III, and 81 days for Stage IV.

For the US-VA dataset, five-year survival was 19.4% for Stage I, 8.8% for Stage II, 3.2% for Stage III, and 1.3% for Stage IV. Median survival was 424 days for Stage I, 330 days for Stage II, 243 days for Stage III, and 91 days for Stage IV.

Ordered for comparison:

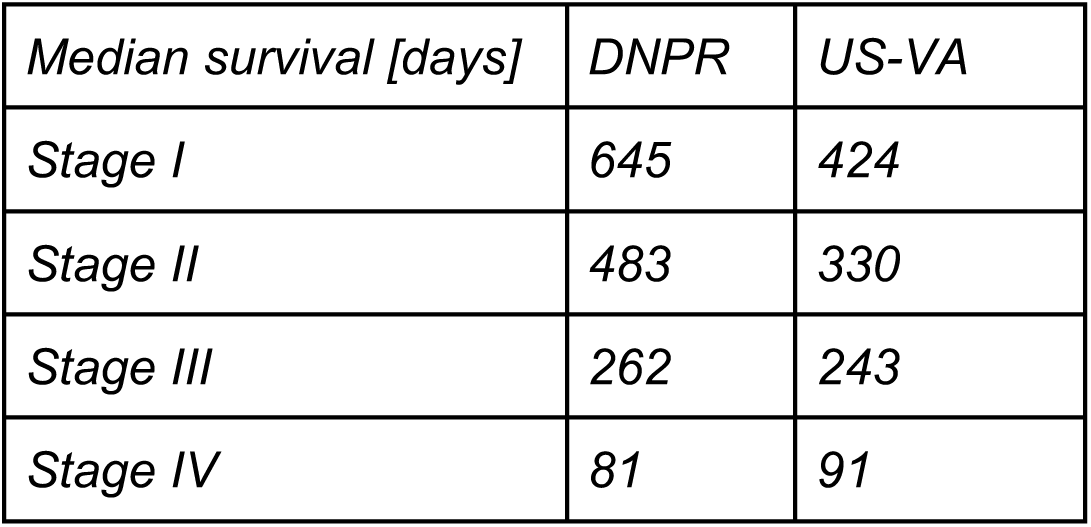

| Median survival [days] | DNPR | US-VA |
| --- | --- | --- |
| Stage I | 645 | 424 |
| Stage II | 483 | 330 |
| Stage III | 262 | 243 |
| Stage IV | 81 | 91 |

The uneven distribution of sex in the VA database may contribute to the differences in survival. The overall similarity of the survival curves in the two datasets adds some confidence to the quality of the data, as selected for the study.

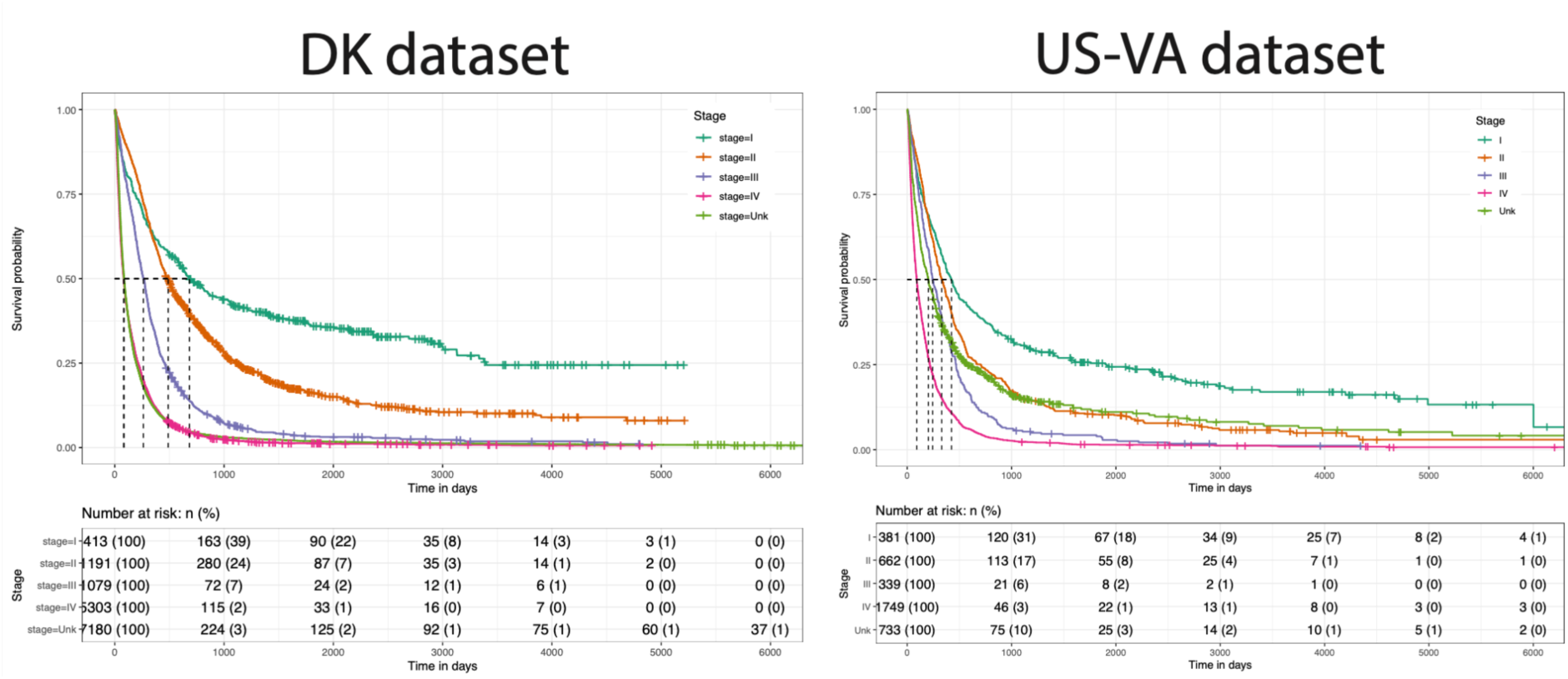

**Table S1.** Description of the patient cohorts used in this study (DK).

| Population Metadata (n=8,110,706 persons) |  |  |
| --- | --- | --- |
| Gender | Male | Female |
| Total Count | 4,030,504 (49.69%) | 4,080,202 (50.31%) |
| Alive | 2,754,152 (33.96%) | 2,827,021 (34.86% ) |
| Dead | 1,276,352 (15.74%) | 1,253,181 (15.45%) |
| After continuity and length filtering | 2,938,248 (36.23%) | 3,239,989 (39.95%) |
| Age at last record [0-10) | 216,329 (2.67%) | 204,774 (2.52%) |
| Age at last record [10-20) | 332,326 (4.10%) | 314,445 (3.88%) |
| Age at last record [20-30) | 322,802 (3.98%) | 298,219 (3.68%) |
| Age at last record [30-40) | 283,200 (3.49%) | 305,470 (3.77%) |
| Age at last record [40-50) | 323,811 (3.99%) | 380,730 (4.69%) |
| Age at last record [50-60) | 368,686 (4.55%) | 419,100 (5.17%) |
| Age at last record [60-70) | 373,220 (4.60%) | 402,625 (4.96%) |
| Age at last record [70-80) | 394,789 (4.87%) | 408,890 (5.04%) |
| Age at last record [80-90) | 258,193 (3.18%) | 342,174 (4.22%) |
| Age at last record [90-100) | 63,470 (0.78%) | 156,154 (1.93%) |
| Age at last record [100, ∞) | 1,422 (0.02%) | 7,398 (0.09%) |

| Pancreatic Cancer Patients (n=23,985) |  |  |
| --- | --- | --- |
|  | Male | Female |
| Total Count | 11,880 (49.53%) | 12,105 50.47% |
| Age at pancreatic cancer diagnosis [0,30) | 13 (0.05%) | 19 (0.08%) |
| Age at pancreatic cancer diagnosis [30-40) | 92 (0.38%) | 93 (0.39%) |
| Age at pancreatic cancer diagnosis [40-50) | 474 (1.98%) | 417 (1.74%) |
| Age at pancreatic cancer diagnosis [50-60) | 1,626 (6.78%) | 1,304 (5.44%) |
| Age at pancreatic cancer diagnosis [60-70) | 3,585 (14.95%) | 2,950 (12.30%) |
| Age at pancreatic cancer diagnosis [70-80) | 4,017 (16.75%) | 4,076 (16.99%) |
| Age at pancreatic cancer diagnosis [80-90) | 1,925 (8.03%) | 2,751 (11.47%) |
| Age at pancreatic cancer diagnosis [90, ∞) | 148 (0.62%) | 495 (2.06%) |

**Table S2.** Hyperparameter search for machine learning models. To comprehensively test different types of neural networks and the corresponding hyperparameters, we conducted a large parameter search for each of the network types. The different types of models include simple feed-forward models (LR, MLP) and more complex models that can take the sequential information of disease ordering into consideration (RNN, GRU and Transformer). The hyperparameters of the best performing model are in **bold**.

|  | Type of ML model |  |  |  |
| --- | --- | --- | --- | --- |
| Hyper-parameters | Bag of words | MLP | GRU | Transformer |
| Dropout | 0 | 0,0.1 | 0,0.1 | **0**, 0.1 |
| Weight decay | 0.001 | 0,0.001 | 0,0.001 | 0, **0.001** |
| Only prior knowledge diseases | False, True | False | False | **False**, True |
| Dimension of hidden layer | - | 32, 128, 256 | 32, 64, 128, 256 | 32, **256** |
| Number of hidden layers | - | 1, 2 | 1, 2, 4 | **1**, 2, 4 |
| Age input | None | None | None, positional embedding | None, **positional embedding** |
| Time input | None | None | None, positional embedding | None, **positional embedding** |
| Number of Heads | - | - | - | 8, **16**, 32 |

**Table S3.**
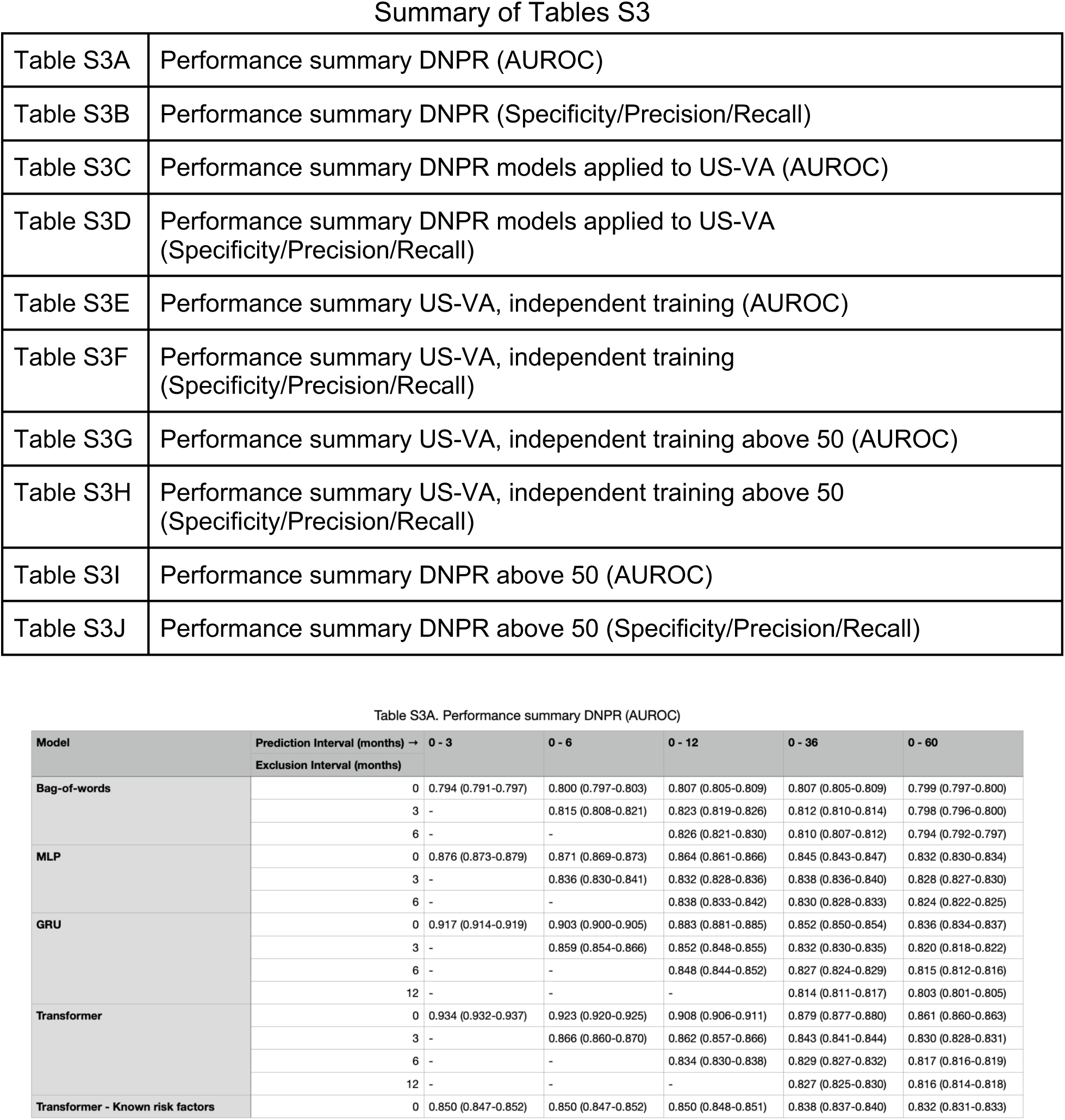
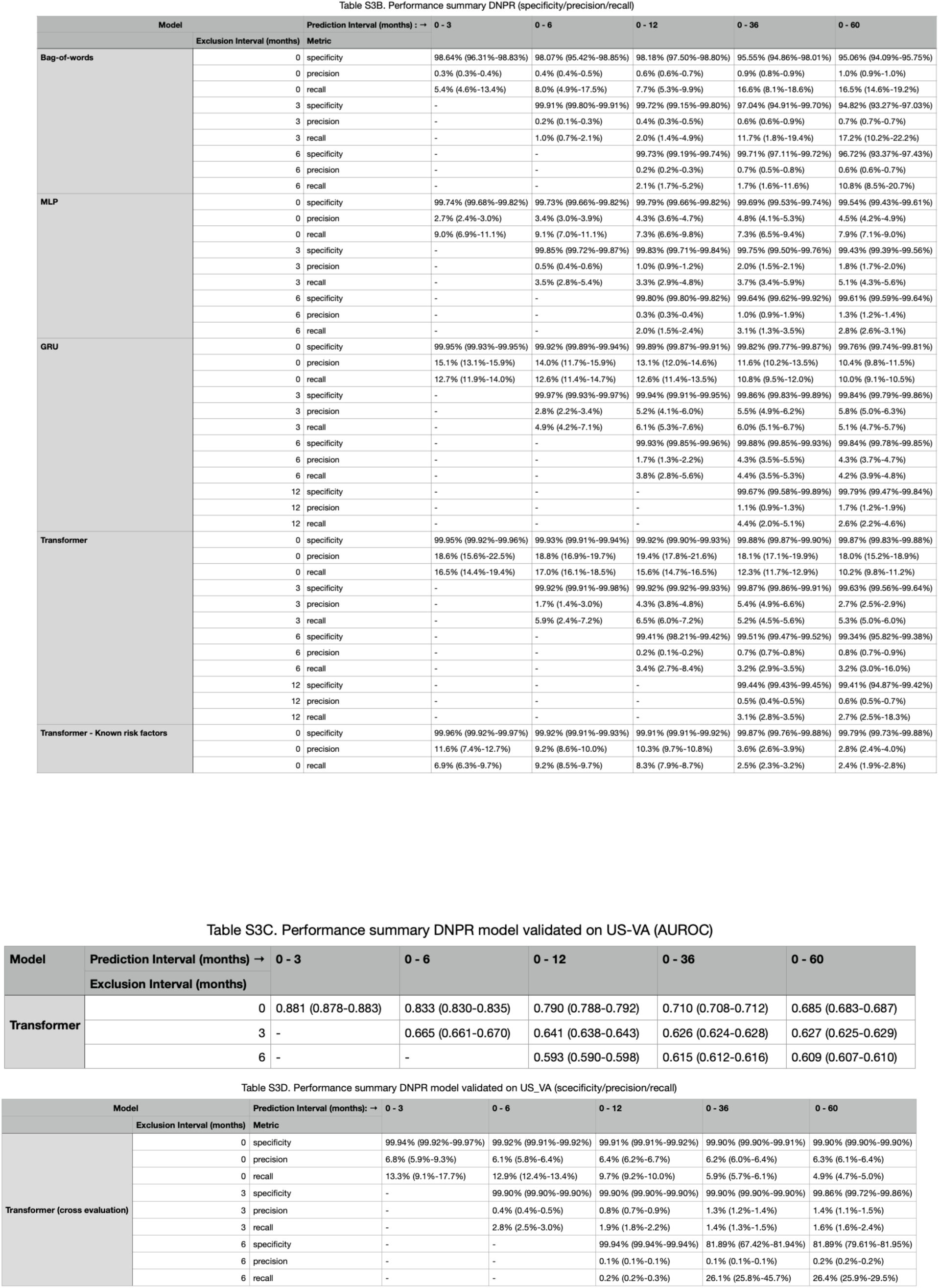
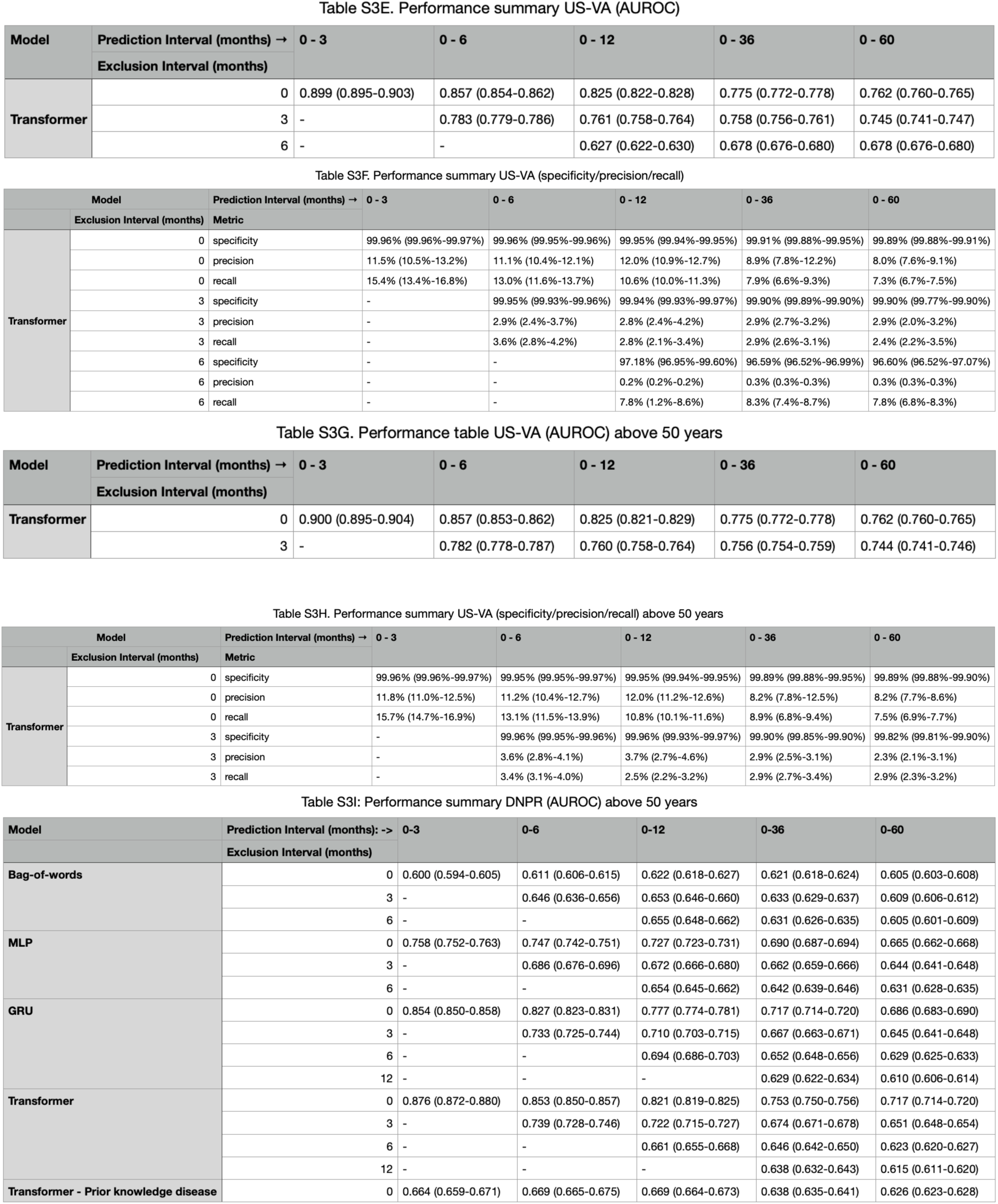
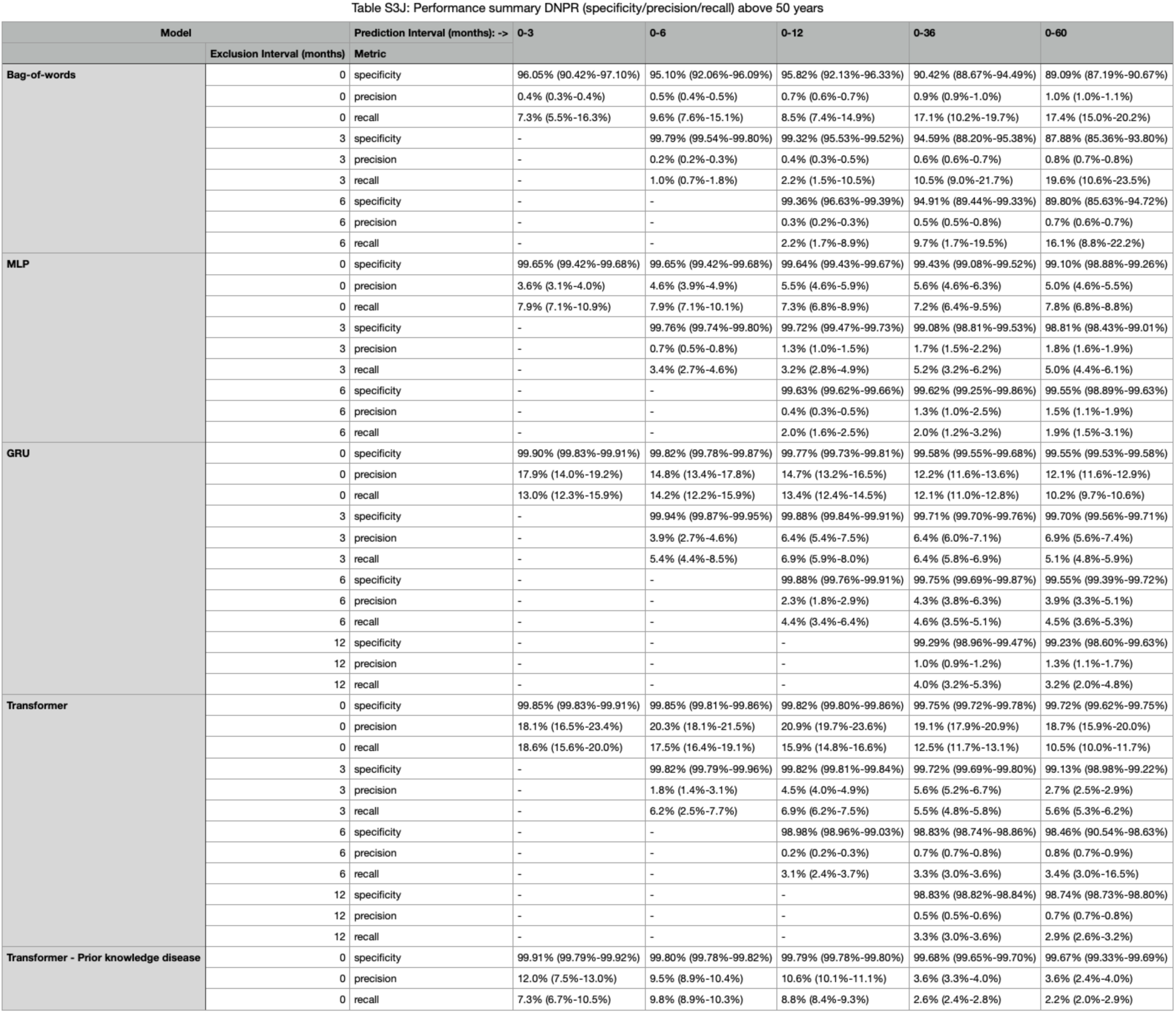
Performance of exclusion experiments - DK dataset and US-VA dataset. A summary of performance of different models trained with different data exclusion intervals for different prediction intervals. In order to estimate the uncertainty of the performance metrics, 95% confidence interval (CI) were computed using 200 resamples (bootstrapping with replacement); these time intervals may be slightly too narrow due to the estimated small number of trajectories from a single patient in a particular sample, but provide a reasonable guide. Specificity, precision, and recall are for the F1-optimal operational point that maximizes the F1 score, which is the harmonic mean of recall and precision (Sasaki 2007).

**Table S4.**
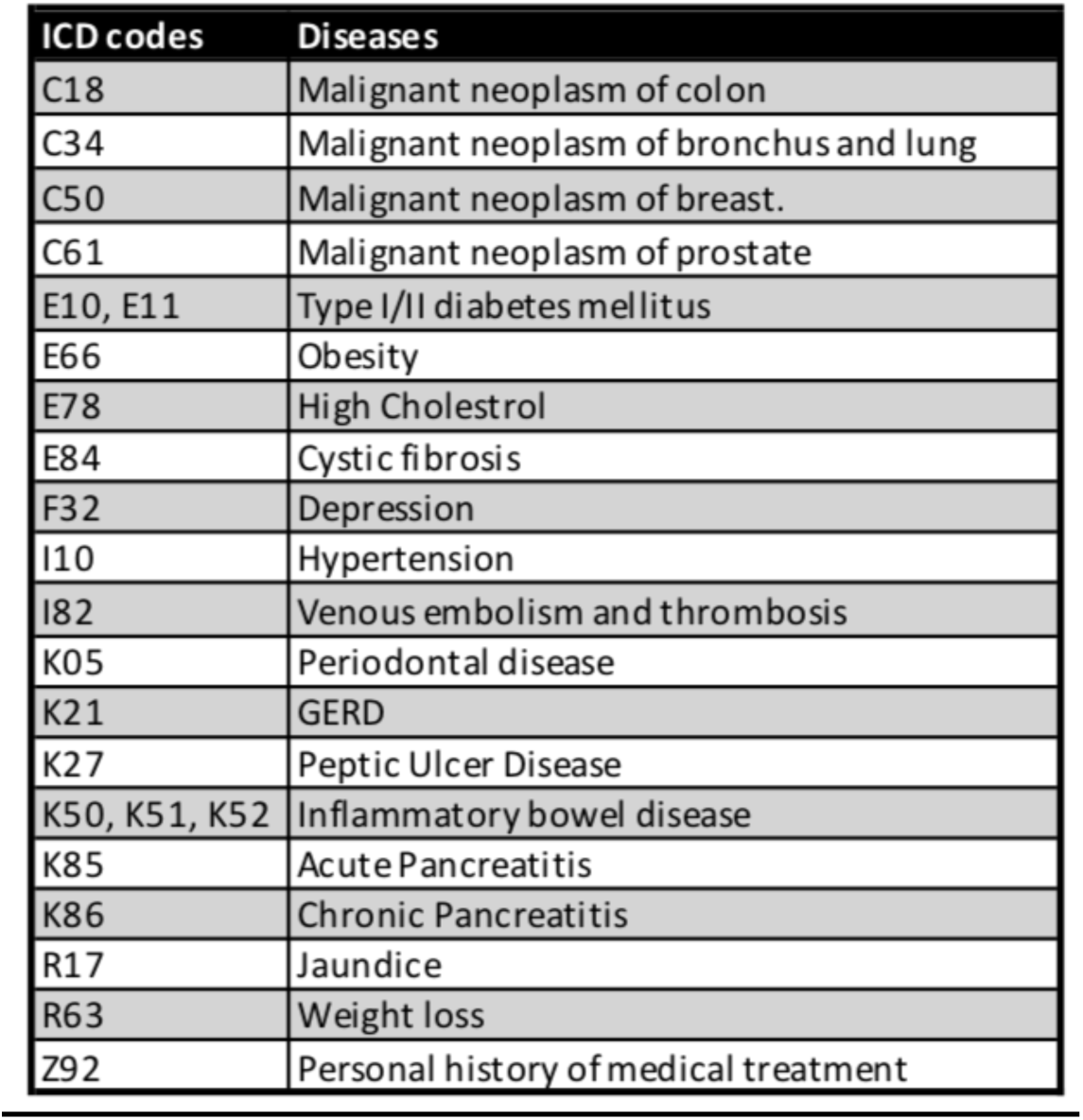
Known risk factor disease codes. A subset of 23 diseases (subset of the 2000 level 3 ICD codes) that have been considered as risk factors for pancreatic cancer (Yuan et al. 2020; Klein 2021) were chosen for the “known risk factor” analysis. Indeed, most of these are flagged by the IG feature extraction method to make a significant contribution to the machine learning prediction of cancer occurrence (**Figure 4**). These risk factors were used to train a separate time-series model ‘Transformer - known risk factors’ for comparison to the model trained on all ICD codes (Figure 3).

**Table S5.**
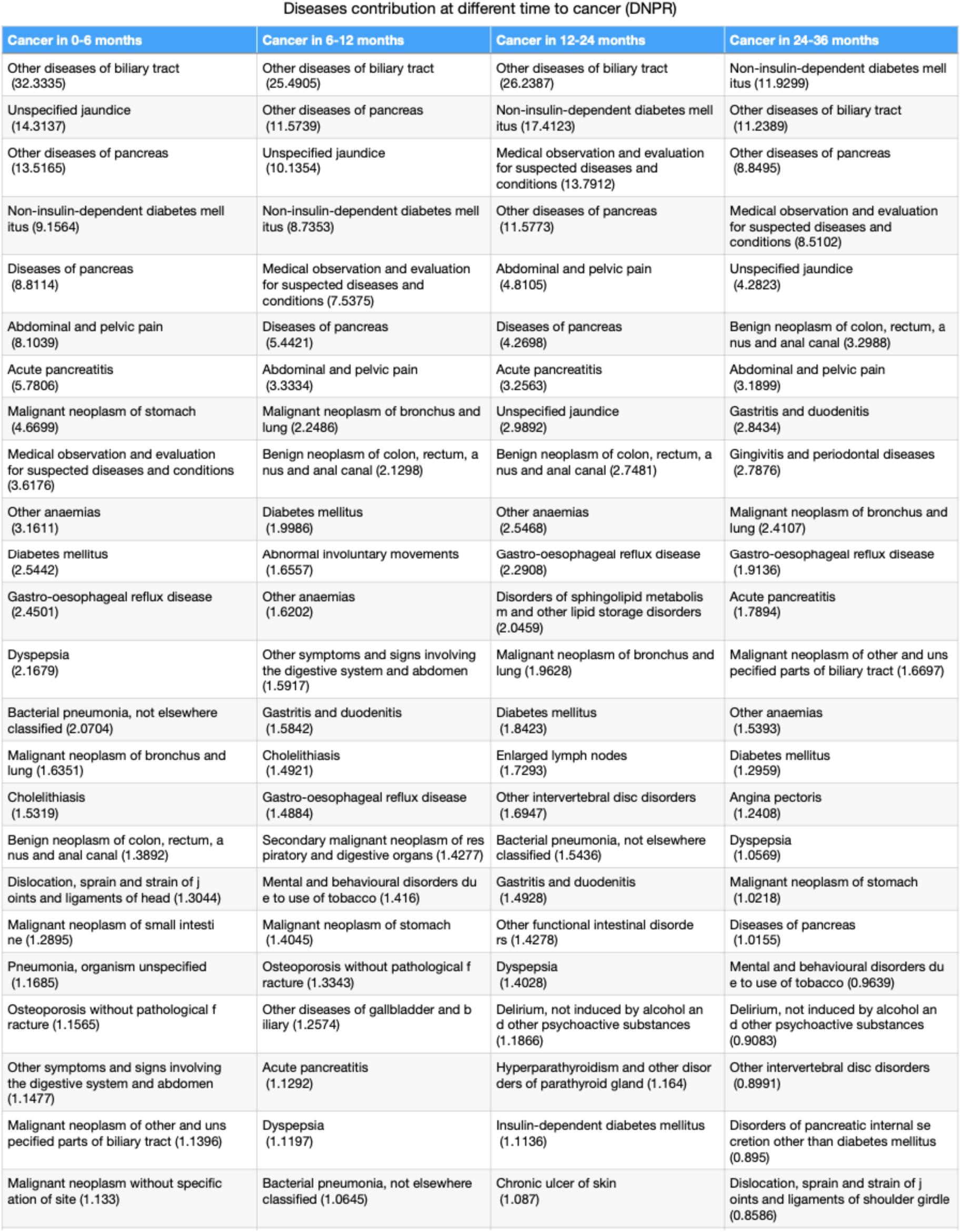
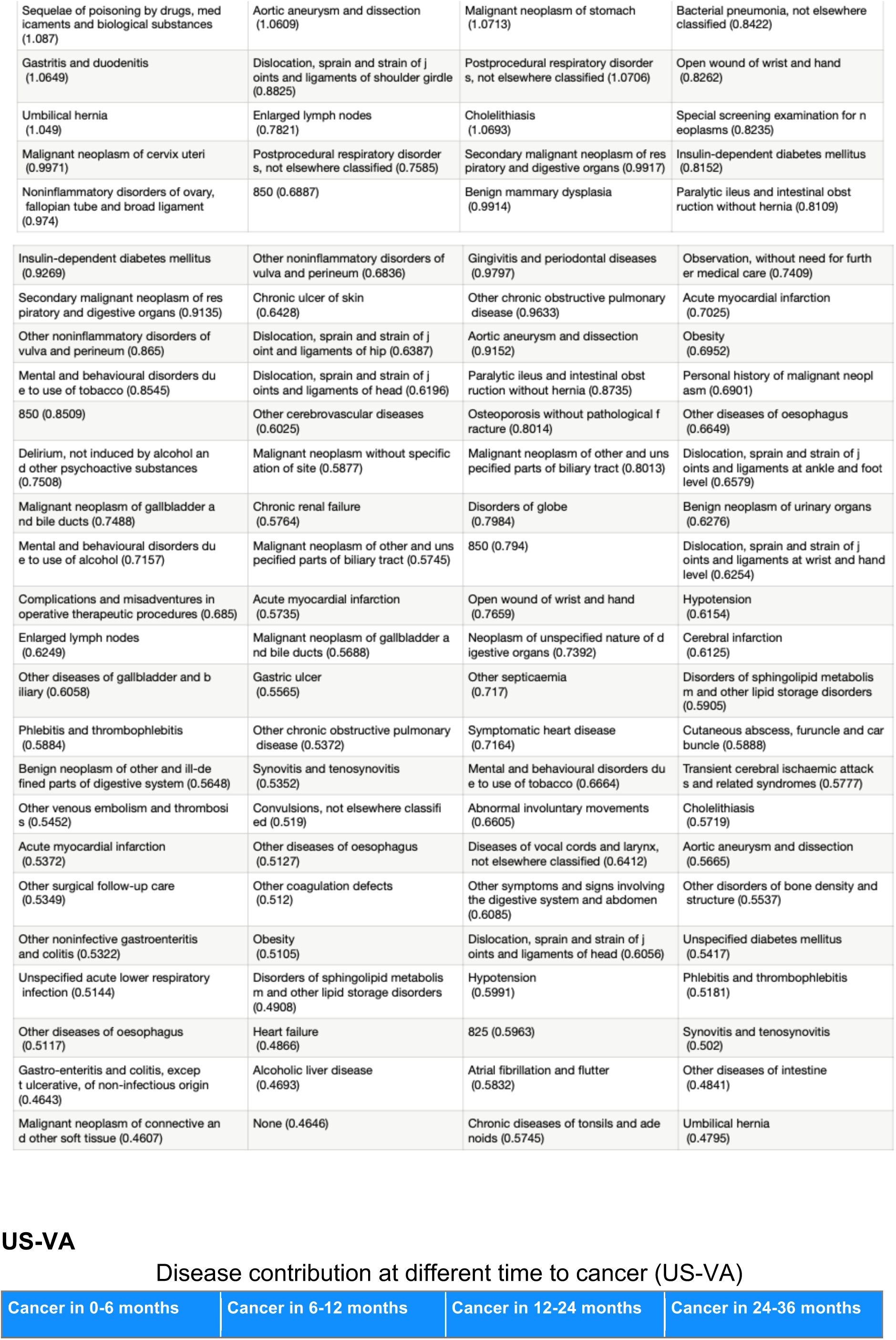
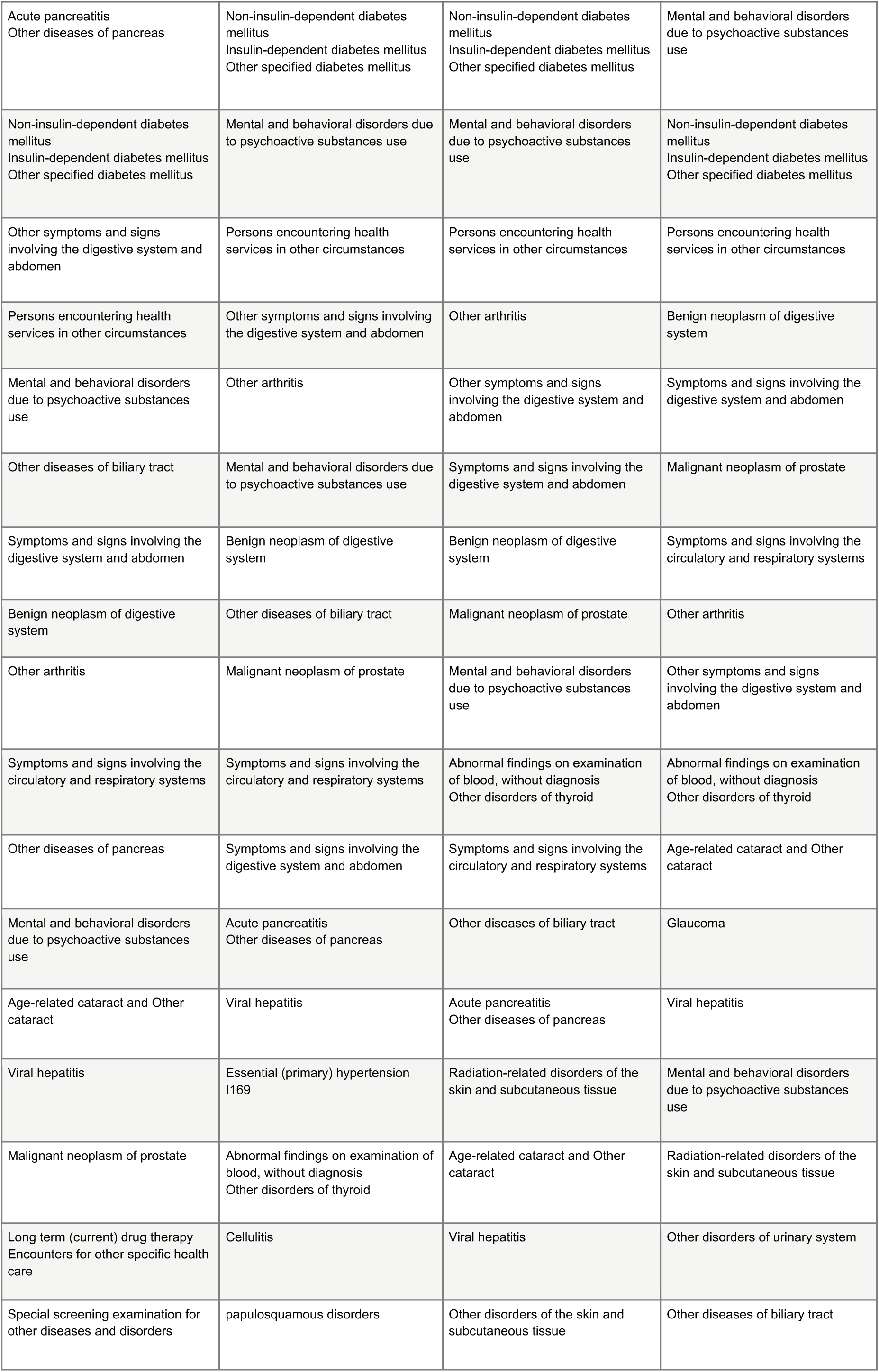
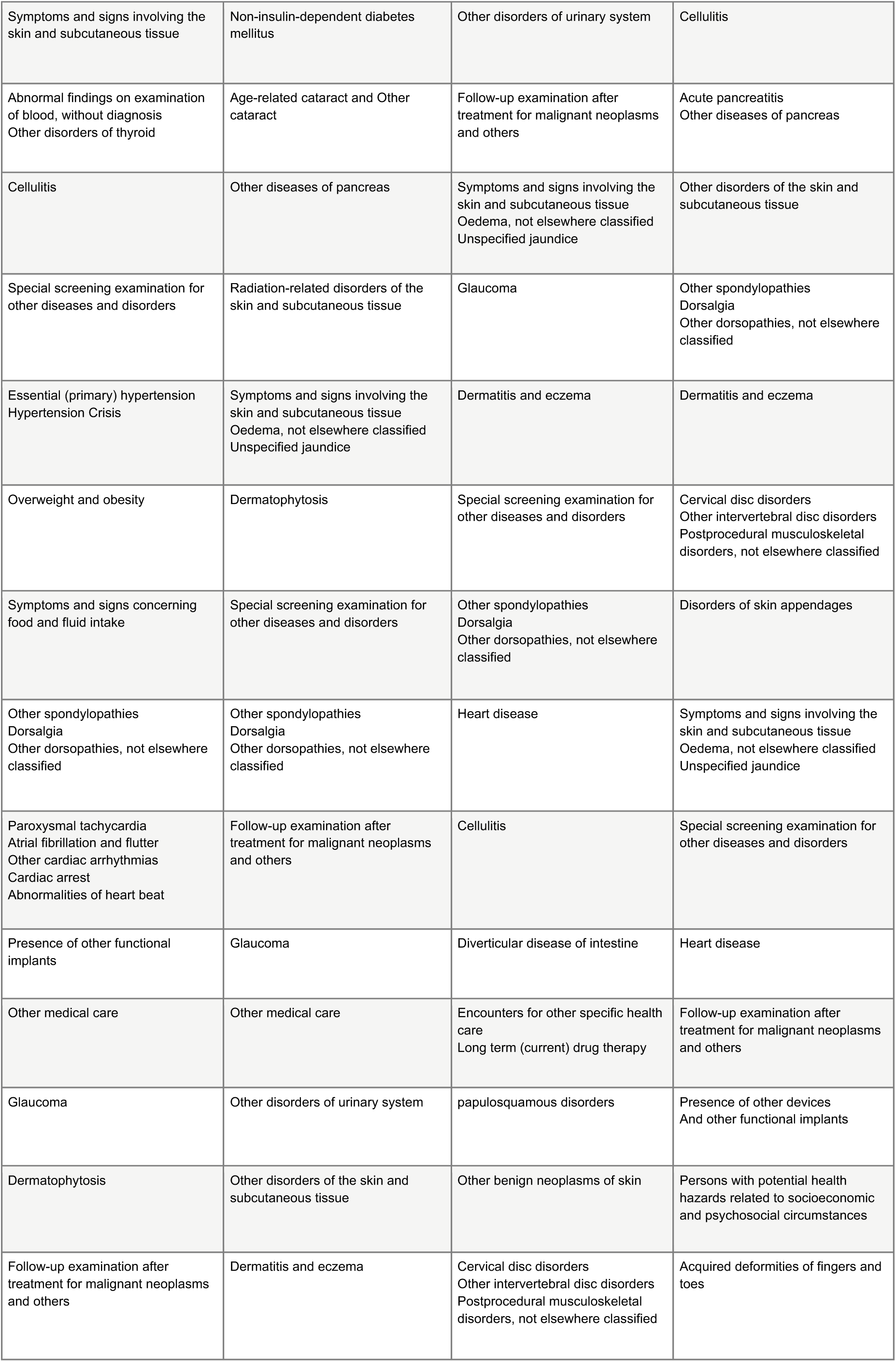
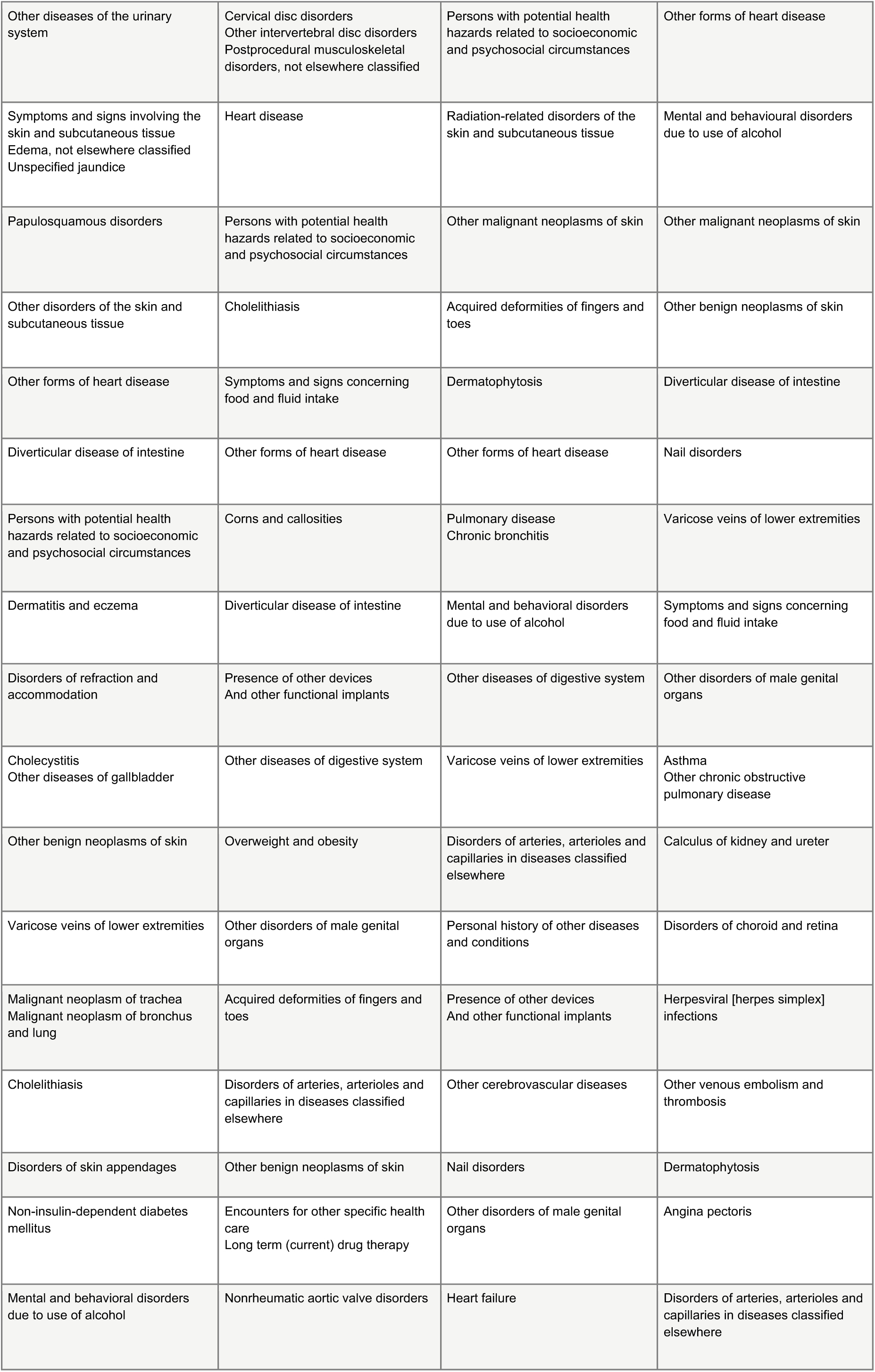
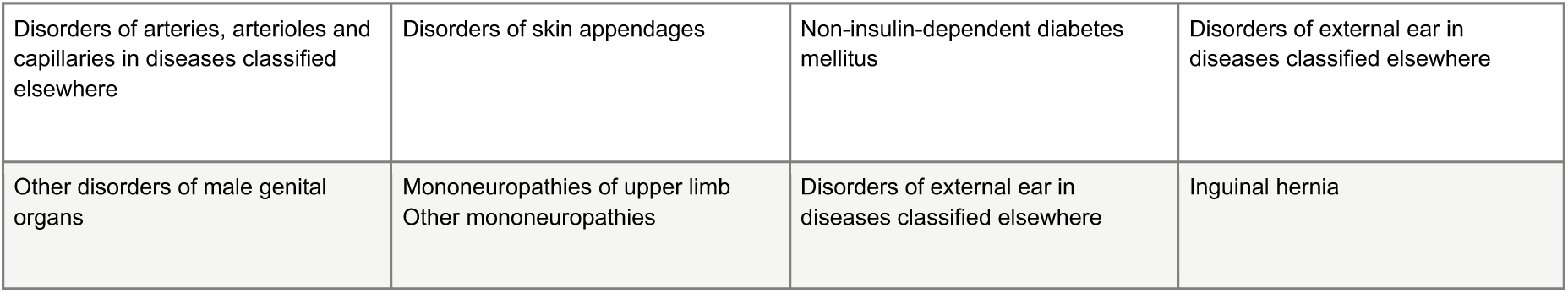
Disease attribution with 3 months data exclusion - Denmark DNPR & US-VA. Ixn order to discover the top diseases that contribute to our model’s risk prediction, we calculated the contribution score for all input features using integrated gradients (IG), an attribution method for neural networks. The IG contribution score (arbitrary units) was calculated for trajectories with cancer occurrence in the time windows 0-6 months, 6-12 months, 12-24 months and 24-36 months for DNPR (**A**) and US-VA (**B**), both with 3 months data exclusion. - A list with features without data exclusion is in **Figure 5**. - We mapped ICD9 diagnosis codes to ICD10 for the US-VA dataset to keep the description comparable, and therefore multiple code descriptions might be listed for a given ICD9 code.

Diseases contribution at different time to cancer (DNPR)
| Cancer in 0-6 months | Cancer in 6-12 months | Cancer in 12-24 months | Cancer in 24-36 months |
| --- | --- | --- | --- |
| Other diseases of biliary tract (32.3335) | Other diseases of biliary tract (25.4905) | Other diseases of biliary tract (26.2387) | Non-insulin-dependent diabetes mellitus (11.9299) |
| Unspecified jaundice (14.3137) | Other diseases of pancreas (11.5739) | Non-insulin-dependent diabetes mellitus (17.4123) | Other diseases of biliary tract (11.2389) |
| Other diseases of pancreas (13.5165) | Unspecified jaundice (10.1354) | Medical observation and evaluation for suspected diseases and conditions (13.7912) | Other diseases of pancreas (8.8495) |
| Non-insulin-dependent diabetes mellitus (9.1564) | Non-insulin-dependent diabetes mellitus (8.7353) | Other diseases of pancreas (11.5773) | Medical observation and evaluation for suspected diseases and conditions (8.5102) |
| Diseases of pancreas (8.8114) | Medical observation and evaluation for suspected diseases and conditions (7.5375) | Abdominal and pelvic pain (4.8105) | Unspecified jaundice (4.2823) |
| Abdominal and pelvic pain (8.1039) | Diseases of pancreas (5.4421) | Diseases of pancreas (4.2698) | Benign neoplasm of colon, rectum, anus and anal canal (3.2988) |
| Acute pancreatitis (5.7806) | Abdominal and pelvic pain (3.3334) | Acute pancreatitis (3.2563) | Abdominal and pelvic pain (3.1899) |
| Malignant neoplasm of stomach (4.6699) | Malignant neoplasm of bronchus and lung (2.2486) | Unspecified jaundice (2.9892) | Gastritis and duodenitis (2.8434) |
| Medical observation and evaluation for suspected diseases and conditions (3.6176) | Benign neoplasm of colon, rectum, anus and anal canal (2.1298) | Benign neoplasm of colon, rectum, anus and anal canal (2.7481) | Gingivitis and periodontal diseases (2.7876) |
| Other anaemias (3.1611) | Diabetes mellitus (1.9986) | Other anaemias (2.5468) | Malignant neoplasm of bronchus and lung (2.4107) |
| Diabetes mellitus (2.5442) | Abnormal involuntary movements (1.6557) | Gastro-oesophageal reflux disease (2.2908) | Gastro-oesophageal reflux disease (1.9136) |
| Gastro-oesophageal reflux disease (2.4501) | Other anaemias (1.6202) | Disorders of sphingolipid metabolism and other lipid storage disorders (2.0459) | Acute pancreatitis (1.7894) |
| Dyspepsia (2.1679) | Other symptoms and signs involving the digestive system and abdomen (1.5917) | Malignant neoplasm of bronchus and lung (1.9628) | Malignant neoplasm of other and unspecified parts of biliary tract (1.6697) |
| Bacterial pneumonia, not elsewhere classified (2.0704) | Gastritis and duodenitis (1.5842) | Diabetes mellitus (1.8423) | Other anaemias (1.5393) |
| Malignant neoplasm of bronchus and lung (1.6351) | Cholelithiasis (1.4921) | Enlarged lymph nodes (1.7293) | Diabetes mellitus (1.2959) |
| Cholelithiasis (1.5319) | Gastro-oesophageal reflux disease (1.4884) | Other intervertebral disc disorders (1.6947) | Angina pectoris (1.2408) |
| Benign neoplasm of colon, rectum, anus and anal canal (1.3892) | Secondary malignant neoplasm of respiratory and digestive organs (1.4277) | Bacterial pneumonia, not elsewhere classified (1.5436) | Dyspepsia (1.0569) |
| Dislocation, sprain and strain of joints and ligaments of head (1.3044) | Mental and behavioural disorders due to use of tobacco (1.416) | Gastritis and duodenitis (1.4928) | Malignant neoplasm of stomach (1.0218) |
| Malignant neoplasm of small intestine (1.2895) | Malignant neoplasm of stomach (1.4045) | Other functional intestinal disorders (1.4278) | Diseases of pancreas (1.0155) |
| Pneumonia, organism unspecified (1.1685) | Osteoporosis without pathological fracture (1.3343) | Dyspepsia (1.4028) | Mental and behavioural disorders due to use of tobacco (0.9639) |
| Osteoporosis without pathological fracture (1.1565) | Other diseases of gallbladder and biliary (1.2574) | Delirium, not induced by alcohol and other psychoactive substances (1.1866) | Delirium, not induced by alcohol and other psychoactive substances (0.9083) |
| Other symptoms and signs involving the digestive system and abdomen (1.1477) | Acute pancreatitis (1.1292) | Hyperparathyroidism and other disorders of parathyroid gland (1.164) | Other intervertebral disc disorders (0.8991) |
| Malignant neoplasm of other and unspecified parts of biliary tract (1.1396) | Dyspepsia (1.1197) | Insulin-dependent diabetes mellitus (1.1136) | Disorders of pancreatic internal secretion other than diabetes mellitus (0.895) |
| Malignant neoplasm without specification of site (1.133) | Bacterial pneumonia, not elsewhere classified (1.0645) | Chronic ulcer of skin (1.087) | Dislocation, sprain and strain of joints and ligaments of shoulder girdle (0.8586) |
| Sequelae of poisoning by drugs, medications and biological substances (1.087) | Aortic aneurysm and dissection (1.0609) | Malignant neoplasm of stomach (1.0713) | Bacterial pneumonia, not elsewhere classified (0.8422) |
| Gastritis and duodenitis (1.0649) | Dislocation, sprain and strain of joints and ligaments of shoulder girdle (0.8825) | Postprocedural respiratory disorders, not elsewhere classified (1.0706) | Open wound of wrist and hand (0.8262) |
| Umbilical hernia (1.049) | Enlarged lymph nodes (0.7821) | Cholelithiasis (1.0693) | Special screening examination for neoplasms (0.8235) |
| Malignant neoplasm of cervix uteri (0.9971) | Postprocedural respiratory disorders, not elsewhere classified (0.7585) | Secondary malignant neoplasm of respiratory and digestive organs (0.9917) | Insulin-dependent diabetes mellitus (0.8152) |
| Noninflammatory disorders of ovary, fallopian tube and broad ligament (0.974) | 850 (0.6887) | Benign mammary dysplasia (0.9914) | Paralytic ileus and intestinal obstruction without hernia (0.8109) |
| Insulin-dependent diabetes mellitus (0.9269) | Other noninflammatory disorders of vulva and perineum (0.6836) | Gingivitis and periodontal diseases (0.9797) | Observation, without need for further medical care (0.7409) |
| Secondary malignant neoplasm of respiratory and digestive organs (0.9135) | Chronic ulcer of skin (0.6428) | Other chronic obstructive pulmonary disease (0.9633) | Acute myocardial infarction (0.7025) |
| Other noninflammatory disorders of vulva and perineum (0.865) | Dislocation, sprain and strain of joints and ligaments of hip (0.6387) | Aortic aneurysm and dissection (0.9152) | Obesity (0.6952) |
| Mental and behavioural disorders due to use of tobacco (0.8545) | Dislocation, sprain and strain of joints and ligaments of head (0.6196) | Paralytic ileus and intestinal obstruction without hernia (0.8735) | Personal history of malignant neoplasm (0.6901) |
| 850 (0.8509) | Other cerebrovascular diseases (0.6025) | Osteoporosis without pathological fracture (0.8014) | Other diseases of oesophagus (0.6649) |
| Delirium, not induced by alcohol and other psychoactive substances (0.7508) | Malignant neoplasm without specification of site (0.5877) | Malignant neoplasm of other and unspecified parts of biliary tract (0.8013) | Dislocation, sprain and strain of joints and ligaments at ankle and foot level (0.6579) |
| Malignant neoplasm of gallbladder and bile ducts (0.7488) | Chronic renal failure (0.5764) | Disorders of globe (0.7984) | Benign neoplasm of urinary organs (0.6276) |
| Mental and behavioural disorders due to use of alcohol (0.7157) | Malignant neoplasm of other and unspecified parts of biliary tract (0.5745) | 850 (0.794) | Dislocation, sprain and strain of joints and ligaments at wrist and hand level (0.6254) |
| Complications and misadventures in operative therapeutic procedures (0.685) | Acute myocardial infarction (0.5735) | Open wound of wrist and hand (0.7659) | Hypotension (0.6154) |
| Enlarged lymph nodes (0.6249) | Malignant neoplasm of gallbladder and bile ducts (0.5688) | Neoplasm of unspecified nature of digestive organs (0.7392) | Cerebral infarction (0.6125) |
| Other diseases of gallbladder and biliary (0.6058) | Gastric ulcer (0.5565) | Other septicaemia (0.717) | Disorders of sphingolipid metabolism and other lipid storage disorders (0.5905) |
| Phlebitis and thrombophlebitis (0.5884) | Other chronic obstructive pulmonary disease (0.5372) | Symptomatic heart disease (0.7164) | Cutaneous abscess, furuncle and carbuncle (0.5888) |
| Benign neoplasm of other and ill-defined parts of digestive system (0.5648) | Synovitis and tenosynovitis (0.5352) | Mental and behavioural disorders due to use of tobacco (0.6664) | Transient cerebral ischaemic attacks and related syndromes (0.5777) |
| Other venous embolism and thromboses (0.5452) | Convulsions, not elsewhere classified (0.519) | Abnormal involuntary movements (0.6605) | Cholelithiasis (0.5719) |
| Acute myocardial infarction (0.5372) | Other diseases of oesophagus (0.5127) | Diseases of vocal cords and larynx, not elsewhere classified (0.6412) | Aortic aneurysm and dissection (0.5665) |
| Other surgical follow-up care (0.5349) | Other coagulation defects (0.512) | Other symptoms and signs involving the digestive system and abdomen (0.6085) | Other disorders of bone density and structure (0.5537) |
| Other noninfective gastroenteritis and colitis (0.5322) | Obesity (0.5105) | Dislocation, sprain and strain of joints and ligaments of head (0.6056) | Unspecified diabetes mellitus (0.5417) |
| Unspecified acute lower respiratory infection (0.5144) | Disorders of sphingolipid metabolism and other lipid storage disorders (0.4908) | Hypotension (0.5991) | Phlebitis and thrombophlebitis (0.5181) |
| Other diseases of oesophagus (0.5117) | Heart failure (0.4866) | 825 (0.5963) | Synovitis and tenosynovitis (0.502) |
| Gastro-enteritis and colitis, except ulcerative, of non-infectious origin (0.4643) | Alcoholic liver disease (0.4693) | Atrial fibrillation and flutter (0.5832) | Other diseases of intestine (0.4841) |
| Malignant neoplasm of connective and other soft tissue (0.4607) | None (0.4646) | Chronic diseases of tonsils and adenoids (0.5745) | Umbilical hernia (0.4795) |

Disease contribution at different time to cancer (US-VA)
| Cancer in 0-6 months | Cancer in 6-12 months | Cancer in 12-24 months | Cancer in 24-36 months |
| --- | --- | --- | --- |
| Acute pancreatitis<br>Other diseases of pancreas | Non-insulin-dependent diabetes mellitus<br>Insulin-dependent diabetes mellitus<br>Other specified diabetes mellitus | Non-insulin-dependent diabetes mellitus<br>Insulin-dependent diabetes mellitus<br>Other specified diabetes mellitus | Mental and behavioral disorders due to psychoactive substances use |
| Non-insulin-dependent diabetes mellitus<br>Insulin-dependent diabetes mellitus<br>Other specified diabetes mellitus | Mental and behavioral disorders due to psychoactive substances use | Mental and behavioral disorders due to psychoactive substances use | Non-insulin-dependent diabetes mellitus<br>Insulin-dependent diabetes mellitus<br>Other specified diabetes mellitus |
| Other symptoms and signs involving the digestive system and abdomen | Persons encountering health services in other circumstances | Persons encountering health services in other circumstances | Persons encountering health services in other circumstances |
| Persons encountering health services in other circumstances | Other symptoms and signs involving the digestive system and abdomen | Other arthritis | Benign neoplasm of digestive system |
| Mental and behavioral disorders due to psychoactive substances use | Other arthritis | Other symptoms and signs involving the digestive system and abdomen | Symptoms and signs involving the digestive system and abdomen |
| Other diseases of biliary tract | Mental and behavioral disorders due to psychoactive substances use | Symptoms and signs involving the digestive system and abdomen | Malignant neoplasm of prostate |
| Symptoms and signs involving the digestive system and abdomen | Benign neoplasm of digestive system | Benign neoplasm of digestive system | Symptoms and signs involving the circulatory and respiratory systems |
| Benign neoplasm of digestive system | Other diseases of biliary tract | Malignant neoplasm of prostate | Other arthritis |
| Other arthritis | Malignant neoplasm of prostate | Mental and behavioral disorders due to psychoactive substances use | Other symptoms and signs involving the digestive system and abdomen |
| Symptoms and signs involving the circulatory and respiratory systems | Symptoms and signs involving the circulatory and respiratory systems | Abnormal findings on examination of blood, without diagnosis<br>Other disorders of thyroid | Abnormal findings on examination of blood, without diagnosis<br>Other disorders of thyroid |
| Other diseases of pancreas | Symptoms and signs involving the digestive system and abdomen | Symptoms and signs involving the circulatory and respiratory systems | Age-related cataract and Other cataract |
| Mental and behavioral disorders due to psychoactive substances use | Acute pancreatitis<br>Other diseases of pancreas | Other diseases of biliary tract | Glaucoma |
| Age-related cataract and Other cataract | Viral hepatitis | Acute pancreatitis<br>Other diseases of pancreas | Viral hepatitis |
| Viral hepatitis | Essential (primary) hypertension I169 | Radiation-related disorders of the skin and subcutaneous tissue | Mental and behavioral disorders due to psychoactive substances use |
| Malignant neoplasm of prostate | Abnormal findings on examination of blood, without diagnosis<br>Other disorders of thyroid | Age-related cataract and Other cataract | Radiation-related disorders of the skin and subcutaneous tissue |
| Long term (current) drug therapy<br>Encounters for other specific health care | Cellulitis | Viral hepatitis | Other disorders of urinary system |
| Special screening examination for other diseases and disorders | papulosquamous disorders | Other disorders of the skin and subcutaneous tissue | Other diseases of biliary tract |
| Symptoms and signs involving the skin and subcutaneous tissue | Non-insulin-dependent diabetes mellitus | Other disorders of urinary system | Cellulitis |
| Abnormal findings on examination of blood, without diagnosis<br>Other disorders of thyroid | Age-related cataract and Other cataract | Follow-up examination after treatment for malignant neoplasms and others | Acute pancreatitis<br>Other diseases of pancreas |
| Cellulitis | Other diseases of pancreas | Symptoms and signs involving the skin and subcutaneous tissue<br>Oedema, not elsewhere classified<br>Unspecified jaundice | Other disorders of the skin and subcutaneous tissue |
| Special screening examination for other diseases and disorders | Radiation-related disorders of the skin and subcutaneous tissue | Glaucoma | Other spondylopathies<br>Dorsalgia<br>Other dorsopathies, not elsewhere classified |
| Essential (primary) hypertension<br>Hypertension Crisis | Symptoms and signs involving the skin and subcutaneous tissue<br>Oedema, not elsewhere classified<br>Unspecified jaundice | Dermatitis and eczema | Dermatitis and eczema |
| Overweight and obesity | Dermatophytosis | Special screening examination for other diseases and disorders | Cervical disc disorders<br>Other intervertebral disc disorders<br>Postprocedural musculoskeletal disorders, not elsewhere classified |
| Symptoms and signs concerning food and fluid intake | Special screening examination for other diseases and disorders | Other spondylopathies<br>Dorsalgia<br>Other dorsopathies, not elsewhere classified | Disorders of skin appendages |
| Other spondylopathies<br>Dorsalgia<br>Other dorsopathies, not elsewhere classified | Other spondylopathies<br>Dorsalgia<br>Other dorsopathies, not elsewhere classified | Heart disease | Symptoms and signs involving the skin and subcutaneous tissue<br>Oedema, not elsewhere classified<br>Unspecified jaundice |
| Paroxysmal tachycardia<br>Atrial fibrillation and flutter<br>Other cardiac arrhythmias<br>Cardiac arrest<br>Abnormalities of heart beat | Follow-up examination after treatment for malignant neoplasms and others | Cellulitis | Special screening examination for other diseases and disorders |
| Presence of other functional implants | Glaucoma | Diverticular disease of intestine | Heart disease |
| Other medical care | Other medical care | Encounters for other specific health care<br>Long term (current) drug therapy | Follow-up examination after treatment for malignant neoplasms and others |
| Glaucoma | Other disorders of urinary system | papulosquamous disorders | Presence of other devices<br>And other functional implants |
| Dermatophytosis | Other disorders of the skin and subcutaneous tissue | Other benign neoplasms of skin | Persons with potential health hazards related to socioeconomic and psychosocial circumstances |
| Follow-up examination after treatment for malignant neoplasms and others | Dermatitis and eczema | Cervical disc disorders<br>Other intervertebral disc disorders<br>Postprocedural musculoskeletal disorders, not elsewhere classified | Acquired deformities of fingers and toes |
| Other diseases of the urinary system | Cervical disc disorders<br>Other intervertebral disc disorders<br>Postprocedural musculoskeletal disorders, not elsewhere classified | Persons with potential health hazards related to socioeconomic and psychosocial circumstances | Other forms of heart disease |
| Symptoms and signs involving the skin and subcutaneous tissue<br>Edema, not elsewhere classified<br>Unspecified jaundice | Heart disease | Radiation-related disorders of the skin and subcutaneous tissue | Mental and behavioural disorders due to use of alcohol |
| Papulosquamous disorders | Persons with potential health hazards related to socioeconomic and psychosocial circumstances | Other malignant neoplasms of skin | Other malignant neoplasms of skin |
| Other disorders of the skin and subcutaneous tissue | Cholelithiasis | Acquired deformities of fingers and toes | Other benign neoplasms of skin |
| Other forms of heart disease | Symptoms and signs concerning food and fluid intake | Dermatophytosis | Diverticular disease of intestine |
| Diverticular disease of intestine | Other forms of heart disease | Other forms of heart disease | Nail disorders |
| Persons with potential health hazards related to socioeconomic and psychosocial circumstances | Corns and callosities | Pulmonary disease<br>Chronic bronchitis | Varicose veins of lower extremities |
| Dermatitis and eczema | Diverticular disease of intestine | Mental and behavioral disorders due to use of alcohol | Symptoms and signs concerning food and fluid intake |
| Disorders of refraction and accommodation | Presence of other devices<br>And other functional implants | Other diseases of digestive system | Other disorders of male genital organs |
| Cholecystitis<br>Other diseases of gallbladder | Other diseases of digestive system | Varicose veins of lower extremities | Asthma<br>Other chronic obstructive pulmonary disease |
| Other benign neoplasms of skin | Overweight and obesity | Disorders of arteries, arterioles and capillaries in diseases classified elsewhere | Calculus of kidney and ureter |
| Varicose veins of lower extremities | Other disorders of male genital organs | Personal history of other diseases and conditions | Disorders of choroid and retina |
| Malignant neoplasm of trachea<br>Malignant neoplasm of bronchus and lung | Acquired deformities of fingers and toes | Presence of other devices<br>And other functional implants | Herpesviral [herpes simplex] infections |
| Cholelithiasis | Disorders of arteries, arterioles and capillaries in diseases classified elsewhere | Other cerebrovascular diseases | Other venous embolism and thrombosis |
| Disorders of skin appendages | Other benign neoplasms of skin | Nail disorders | Dermatophytosis |
| Non-insulin-dependent diabetes mellitus | Encounters for other specific health care<br>Long term (current) drug therapy | Other disorders of male genital organs | Angina pectoris |
| Mental and behavioral disorders due to use of alcohol | Nonrheumatic aortic valve disorders | Heart failure | Disorders of arteries, arterioles and capillaries in diseases classified elsewhere |
| Disorders of arteries, arterioles and capillaries in diseases classified elsewhere | Disorders of skin appendages | Non-insulin-dependent diabetes mellitus | Disorders of external ear in diseases classified elsewhere |
| Other disorders of male genital organs | Mononeuropathies of upper limb<br>Other mononeuropathies | Disorders of external ear in diseases classified elsewhere | Inguinal hernia |

**Figure S2.**
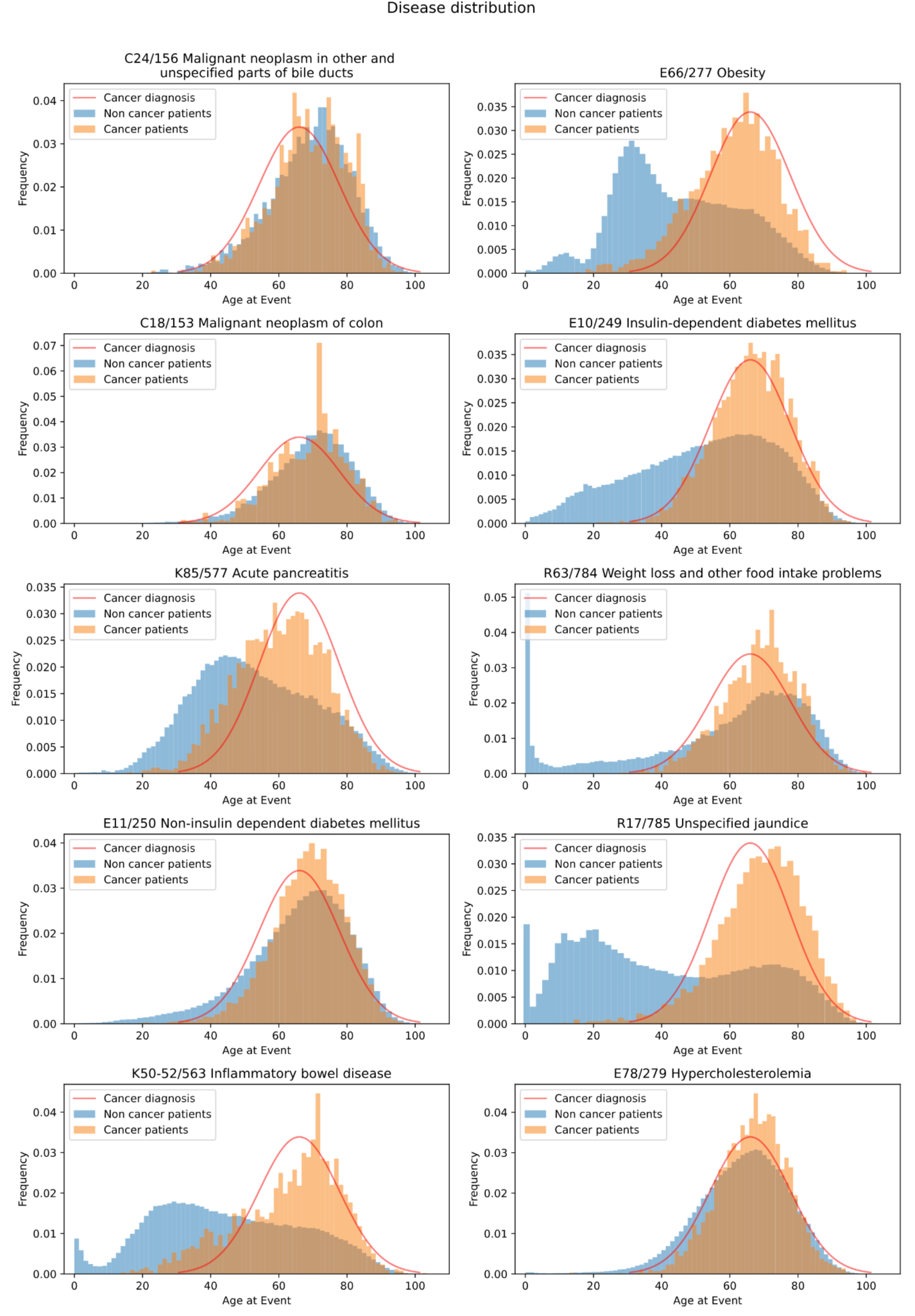
Distribution of disease codes as a function of age in the DNPR (Denmark) database. Distribution of disease codes for a representative subset of diseases known to contribute to the risk of pancreatic cancer, as a fraction of all pancreatic cancer patients (orange) and all non-cancer patients (blue). The similarity of the distributions for some of these diseases with the distribution of occurrence of pancreatic cancer (red line, Gaussian fit to cancer diagnosis data) is consistent with either a direct or indirect contribution to cancer risk -but not taken as evidence in this work. The disease codes are ICD-10/ICD-8.

**Figure S3.**
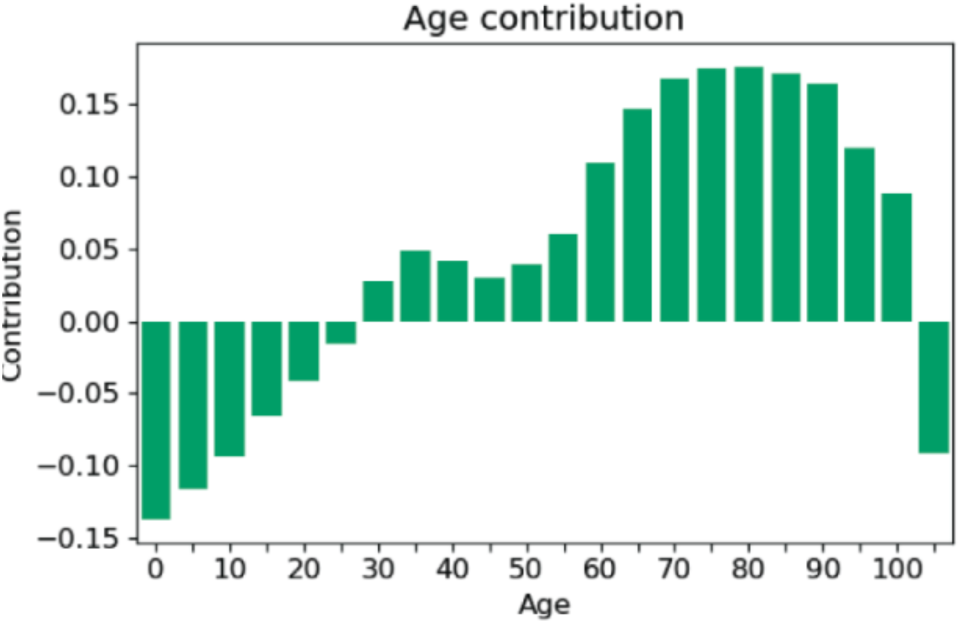
Age as a contributing factor. The integrated gradient method was used to extract the contribution (arbitrary units) of patient age to the prediction at the time of assessment. This confirmed that the positive contribution to risk rises strongly from age 50. As for the disease contributions, the age contribution was calculated in relation to the 3 year (after the time of assessment/prediction) cancer risk.

### Result S1: Draft economic considerations for the design of clinical screening trial

We propose a toy estimate of a practical scenario for a screening trial, taking into account typically available real-world data, the accuracy of prediction on such data, the estimated cost of a screening trial, the cost of clinical screening methods and the overall potential benefit of treatment.

The detailed design of a prediction-surveillance program, to be explored in a clinical setting, depends on the organization of a particular health care system. In a ‘walk in’ scenario, in approximate analogy to colonoscopic screening for colorectal cancer, patients older than, e.g., age 50 would be invited for assessment of their risk by the prediction tool on a yearly basis and, if identified as high-risk, offered a series of follow-up visits with extensive clinical testing. In a ‘national system’ scenario, possible in centralized health systems with location-independent centralized aggregation of electronic health records, risk assessment could be done on an ongoing basis, possibly for each patient whenever a new disease event occurs. If a high-risk prediction is triggered, the responsible physician would receive an alert. With a diversity of scenarios, it is reasonable to propose a clinical prediction-surveillance program tailored to the health system in a particular country.

To illustrate the economic benefits of such a screening and to stimulate discussion regarding the optimal design of a prediction-surveillance program, we have made a first-order-estimate for a clinical screening trial of 10,000 people using the best model (the transformer model). For simplicity, we have made no assumptions regarding age distribution. Here is a simple economic model.

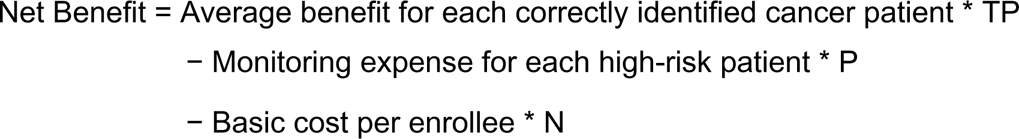

where the screening cohort is N=10,000 and TP is the number of true positives, i.e., the number of correctly identified high-risk patients, and P is the number of actual positive patients, which we estimated using cancer incidence of the DNPR dataset. In our cost-benefit estimate, we arbitrarily set the screening trial cost at $200 per enrollee, the additional monitoring expense for a patient predicted at high risk by screening at $10,000 and the extra cost saved for advanced treatment for each monitored patient at $200,000, averaged over those in which cancer is detected (savings in excess of $200,000) and those in which it is not detected (no savings).

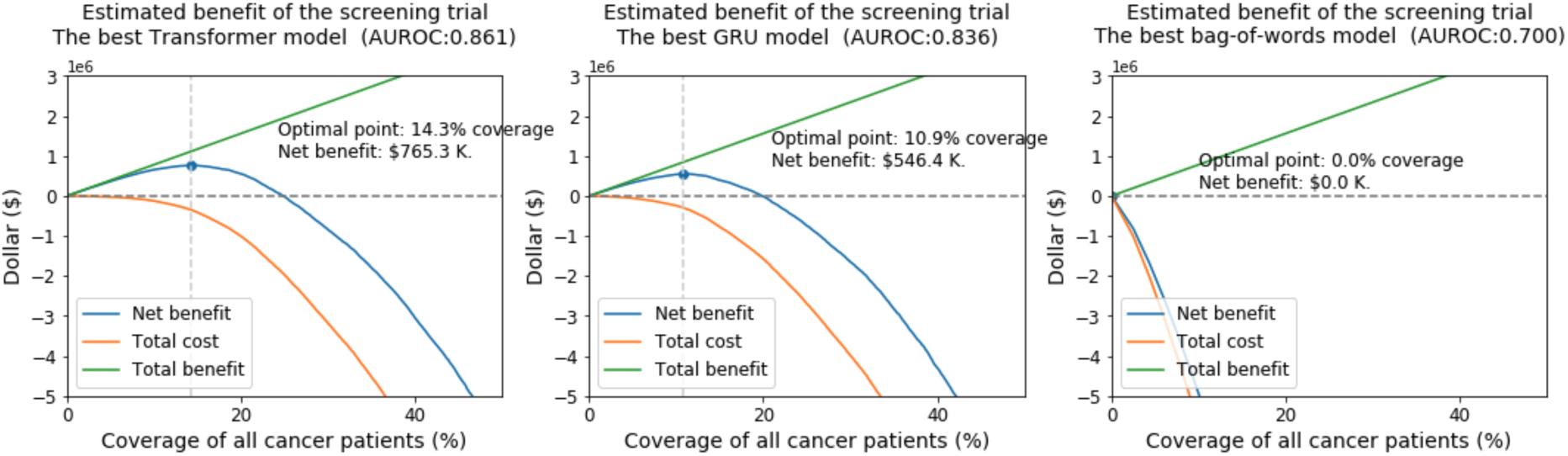

### Estimate of financial benefits for different models

We analyzed each possible operational point and calculated the corresponding cost and benefit, using ballpark estimates. We plotted the net benefits as a function of coverage of cancer patients, i.e. recall or sensitivity. Covering more cancer patients plausibly leads to a larger total benefit, but the total cost also increases. The optimal point is picked for maximal net benefit.

An optimal decision threshold has to balance the cost of assessment and testing against the potential financial benefit for reducing treatment cost. Using this simplified model, we estimated the net benefits of different models with all possible operational points. Such a screening trial for 10,000 people would have $760,000 net benefit by choosing the balance between true and false positives such that the net benefit is optimal. This corresponds to a precision of 14.0% and a specificity of 99.7%. In contrast, a less good model GRU would have $540K net benefits but a bag-of-words model (baseline) would have no net benefits for any operational point because of the low incidence of pancreatic cancer.

The proposed concrete but hypothetical design of a screening trial is intended to guide the debate and ultimate decisions regarding implementation with clinicians and healthcare professionals. However, this calculation is based on roughly estimated numbers and does not reflect real-world cost analysis. Nor does this economic model reflect the non-monetary benefits to patients’ quality of life, which should be the dominant factor in the design of prediction-surveillance and early intervention programs. In a real-world scenario, clinicians and payers in a particular health system have the opportunity to optimize the design of such trials with realistic cost-benefit parameters, as well as consideration of communication ethics and the non-financial aspects of patient benefit.

A key challenge for future realistic economic estimates is the mapping between ICD (diagnosis) codes to CPT (billing) codes that are used for expense calculations and reimbursements. In addition, in the US, there is substantial geographical variability in reimbursement even for the same CPT/billing codes.

